# Selective translation of epigenetic modifiers drives the developmental clock of neural stem cells

**DOI:** 10.1101/2020.10.08.330852

**Authors:** Quan Wu, Yuichi Shichino, Takaya Abe, Taeko Suetsugu, Ayaka Omori, Hiroshi Kiyonari, Shintaro Iwasaki, Fumio Matsuzaki

## Abstract

The cerebral cortex is formed by diverse neurons generated sequentially from neural stem cells (NSCs). A clock mechanism has been suggested to underlie the temporal progression of NSCs, which is mainly defined by the transcriptome and the epigenetic state. However, what drives such a developmental clock remains elusive. We show that translational control of histone H3 trimethylation at Lys27 (H3K27me3) modifiers is part of this clock. We found that depletion of Fbl, an rRNA methyltransferase, reduces translation of both the Ezh2 methyltransferase and Kdm6b demethylase of H3K27me3 and delays progression of the NSC state. These defects are phenocopied by simultaneous inhibition of H3K27me3 methyltransferase and demethylase, indicating the role of Fbl in the genome-wide H3K27me3 pattern. Fbl selectively enhances the translation of H3K27me3 modifiers via a cap-independent mechanism. We thus propose that Fbl drives the intrinsic clock through the translational enhancement of H3K27me3 modifiers that predominantly define the NSC state.

## Main

How the developmental schedule is shared by all individuals in a given species of animals is a fundamental question in developmental biology. One fascinating hypothesis is the presence of a developmental clock that counts time for the developmental program, and there are several potential mechanisms that could work in this manner. During somitogenesis, a clock consisting of a complex gene regulatory network generates oscillation and regulates segmentation in a defined time ^1^. The cell cycle is also an oscillator that counts time to initiate transcription of the zygotic genome during the midblastula transition of *Xenopus* and *Drosophila* ^2,3^. In oligodendrocyte precursors of rat optic nerve, an hourglass type clock was observed: an accumulated amount of a CDK inhibitor p27/kip1 during proliferation of oligodendrocyte precursors determines the timing for their differentiation ^4^. Moreover, an epigenetic clock based on DNA methylation is a promising predictor of biologic age ^5^.

Sequential generation of diverse neurons from a small population of neural stem cells (NSCs) in a highly orchestrated order in the mammalian cerebral cortex may also be controlled by a developmental clock. Following proliferation, NSCs divide asymmetrically to produce one stem cell and either a neuron or intermediate progenitor, the majority of which divide once before terminal differentiation ^6,7^. As neurogenesis proceeds, a shift in NSC gene expression, or identity, occurs; thus, NSC identity is temporally patterned and initiates production of a diverse array of neuronal progeny. NSCs initially produce deep-layer (early-born) neurons, followed by upper-layer (late-born) neurons, and finally, glia ^8,9^.

Temporal patterning of NSC identity is widely observed among species, from the central brain of *Drosophila*, to the mammalian retina, implying the existence of a conserved strategy for neuronal production ^8,10^. In *Drosophila*, key temporal determinant genes have been identified, and epigenetic mechanisms are involved in regulating these genes. For example, NSC expression of the *Hunchback* gene is epigenetically restricted by the relocation of the *Hunchback* locus into a repressive subnuclear compartment ^11^. In the mammalian cortex, a set of temporal genes concordant with temporal identity progression have been identified ^10,12^. Moreover, the perturbation of epigenetic modifier function has been shown to interrupt temporal patterning in mammalian NSCs. For example, perturbation of polycomb repressive complex 2, an epigenetic modifier of histone H3 trimethylation at Lys27 (H3K27me3), compromises temporal shifts in NSC identity, leading to disordered production of NSC progeny cells ^12-15^. Thus, the precise temporal pattern of NSC gene expression largely depends on the precise control of temporal genome-wide epigenetic modifications. However, it remains unclear whether genome-wide epigenetic modification can work as a developmental clock to predict the temporal identity of NSCs, and if so, what factors drive this clock.

## Results

### Dynamics of genome-wide H3K4me3 and H3K27me3 distribution during temporal patterning of NSCs

We first investigated temporal identity changes of NSCs by performing transcriptome analysis at the single-cell level from embryonic (E) day 11 (which is mostly proliferative or at the early neurogenic stage) to E14 (producing later-born neurons at the mid-neurogenic stage) (Fig. 1a). We then constructed a continuous trajectory of all cells including NSCs, neural progenitors, and neurons by a force-directed k-nearest neighbor graph using SPRING ^16^ and interpreted the cell clusters based on known markers (Fig. 1b and Extended Data Fig. 1a). Consistent with a previous model, NSCs gradually change their transcriptome to produce different types of neurons ^10^.We extracted early- and late-onset genes which showed higher expression level in E11 and E14, respectively (Fig. 1c as examples; Extended Data Table 1a for the list).

**Figure 1.**
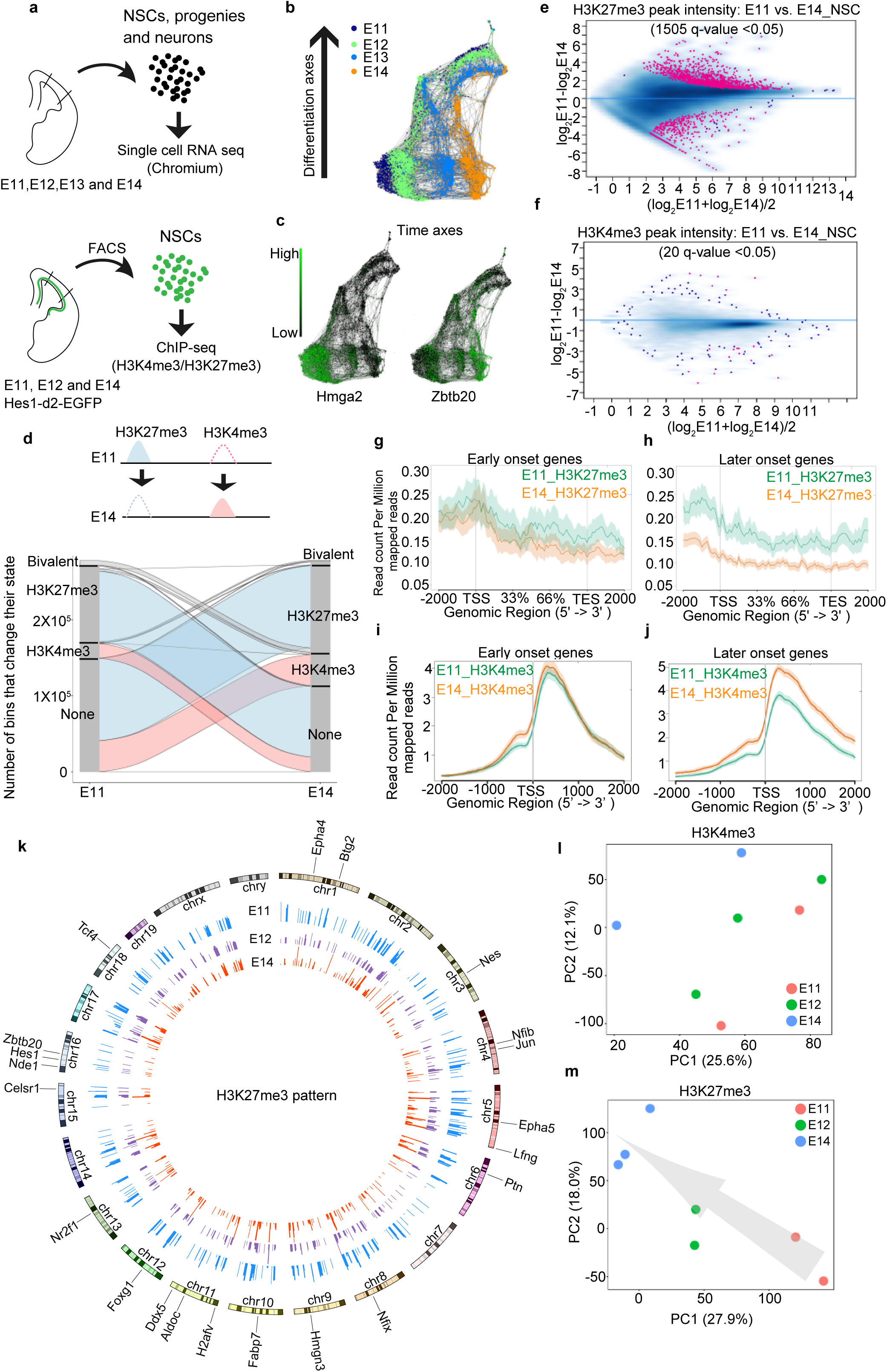
Genome-wide H3K4me3 and H3K27me3 modification change during temporal patterning of NSCs. **a**, Experiment design for single cell (sc) RNA-seq and ChIP-seq. Cell number in scRNA were E11 n=829, E12 n= 2457, E13 n=1566 and E14 n=1293. **b,c**, SPRING graph of single cells coloured by different stages (b), and expression patterns of an early- and a late-onset gene, *Hmga2* and *Zbtb20*, respectively (c). **d**, H3K27me3 and H3K4me3 change of isolated Hes1+ NSCs from E11 to E14. Lines represent 200-bp chromosome regions. **e,f**, Intensity comparison of H3K27me3 (e) and H3K4me3 (f) peaks between E11 and E14 NSCs, showing that H3K27me3 changed more dramatically than H3K4me3. **g-j**, Read-density profiling of H3K27m3 (g, h) and H3K4m3 (i, j) at early-onset (g, i) and late-onset genes (h, j) in wild type E11 and E14 NSCs. **k**, Circos plot showing H3K27me3 pattern change from E11 to E14. Several late-onset genes were highlighted and de-repression of these genes can be observed at E14. **l,m**, Principal component analysis (PCA) of H3K27me3 and H3K4me3 peaks, showing H3K27me3 samples can represent developmental time. Arrow indicates temporal axes.

To investigate the presence of an epigenetic clock that can predict temporal identity of NSCs, we first focused on two major histone modifications: histone H3 trimethylation at Lys4 (H3K4me3) and H3K27me3, which is an active and repressive marker for gene expression, respectively, because temporal identity progression is accompanied with dramatic transcriptome change (Fig. 1b). We performed H3K4me3 and H3K27me3 chromatin immunoprecipitation and sequencing (ChIP-seq) using fluorescence-activated cell sorting (FACS)-sorted NSCs from *Hes1-d2-EGFP* reporter mice, in which d2-GFP is expressed under the control of the promoter of an established NCS marker: *Hes1* ^17^, from E11, E12 and E14 (Fig. 1a). We then analyzed how histone modification changes associated with the temporal identity change of NSCs. We first analyzed the qualities of ChIP-seq data according to several criteria (Extended Data Fig. 1b-f). We subsequently classified each 200bp chromosome region at each stage into one of four states based on two histone modification profiles: H3K4me3-only, H3K27me3-only, bivalent, and no-marker. The majority of genomic regions gained or lost H3K27me3 modification when shifting from E11 to E14. In contrast, changes in ‘H3K4me3-only’ and ‘bivalent’ regions were restricted to relatively small genomic regions (Fig. 1d). We extracted 1505 and 20 sites showing significantly differential intensities (q-value < 0.05) of H3K27me3 and H3K4me3 peaks, respectively, between E11 and E14 NSCs (Fig. 1e,f; and Extended Data Table 1b). Then, we focused on changes in those early- and late-onset genes as described above. Whereas the abundance of H3K27me3 peaks on early-onset genes did not show a clear difference between E11 and E14, it was decreased at E14 compared to E11 in late-onset genes (Fig. 1g,h), implying that H3K27me3 represses the expression of late-onset genes in E11 NSCs (Fig. 1k for some examples). On the other hand, the intensity of H3K4me3 peaks around the transcription start sites of early-onset genes was not drastically changed between E11 and E14 NSCs (Fig. 1i). In contrast, these peaks for late-onset genes increased from E11 to E14, reflecting the higher expression of these genes at E14 (Fig. 1j). Thus, we concluded that dynamic changes in H3K27me3 and H3K4me3 deposition on later-onset genes are associated with the temporal progression of NSCs.

### Global H3K27me3 pattern can predict the developmental time of NSCs

To test whether genome-wide H3K4me3 and H3K27me3 patterns can predict developmental time of NSCs, we performed principle component analysis (PCA) based on the intensity of individual H3K27me3 and H3K4me3 peaks (methods in detail). H3K27me3 samples from different stages can be clearly separated and were deposited along a time axes, while H3K4me3 samples were less distinguished especially for samples from E11 and E12 (Fig. 1l,m). As H3K27me3 patterns are highly related to the developmental stage, we conclude that H3K27me3 patterns within the genome can be considered as a part of the developmental clock in NSCs (Fig. 1k).

### Identification of Fbl as a key regulator of temporal patterning

We then asked what factors promote the developmental clock and lead to temporal identity transition of NSCs. We expected the presence of genes showing monotonic changes of expression among the factors promoting the temporal pattern. Therefore, we compared our single cell transcriptome data from E11 and E14 NSCs ^10^. Using weighted correlation network analysis (WGCNA) ^18^, we identified a gene module with higher expression in E11 than in E14 NSCs (brown module in Extended Data Fig. 2a) that was highly enriched in genes whose products are located in nuclear regions essential for rDNA transcription and pre-rRNA processing, such as nucleolar part and fibrillar center (Fig. 2a,b; and Extended Data Table 2). Among these genes, *Fbl* (also known as *Fibrillarin*) is of particular interest. Fbl was initially reported as an rRNA methyltransferase for 2′-O-methylation and plays an essential role in development and disease ^19-22^. Though Fbl is regarded as essential for the translational regulation of some mRNAs ^23^, its role and underlying mechanisms in mouse brain development are unclear. To address these issues, we first investigated the expression pattern of Fbl. Ubiquitous expression of Fbl in both NSCs and neurons at E11 and E14 was observed by immunostaining (Fig. 2c and Extended Data Fig. 2b). Using western blot, we found higher Fbl protein levels in E11 than E14 FACS-isolated NSCs from *Hes1-d2-EGFP* reporter mice (Extended Data Fig. 2c,d).

**Figure 2.**
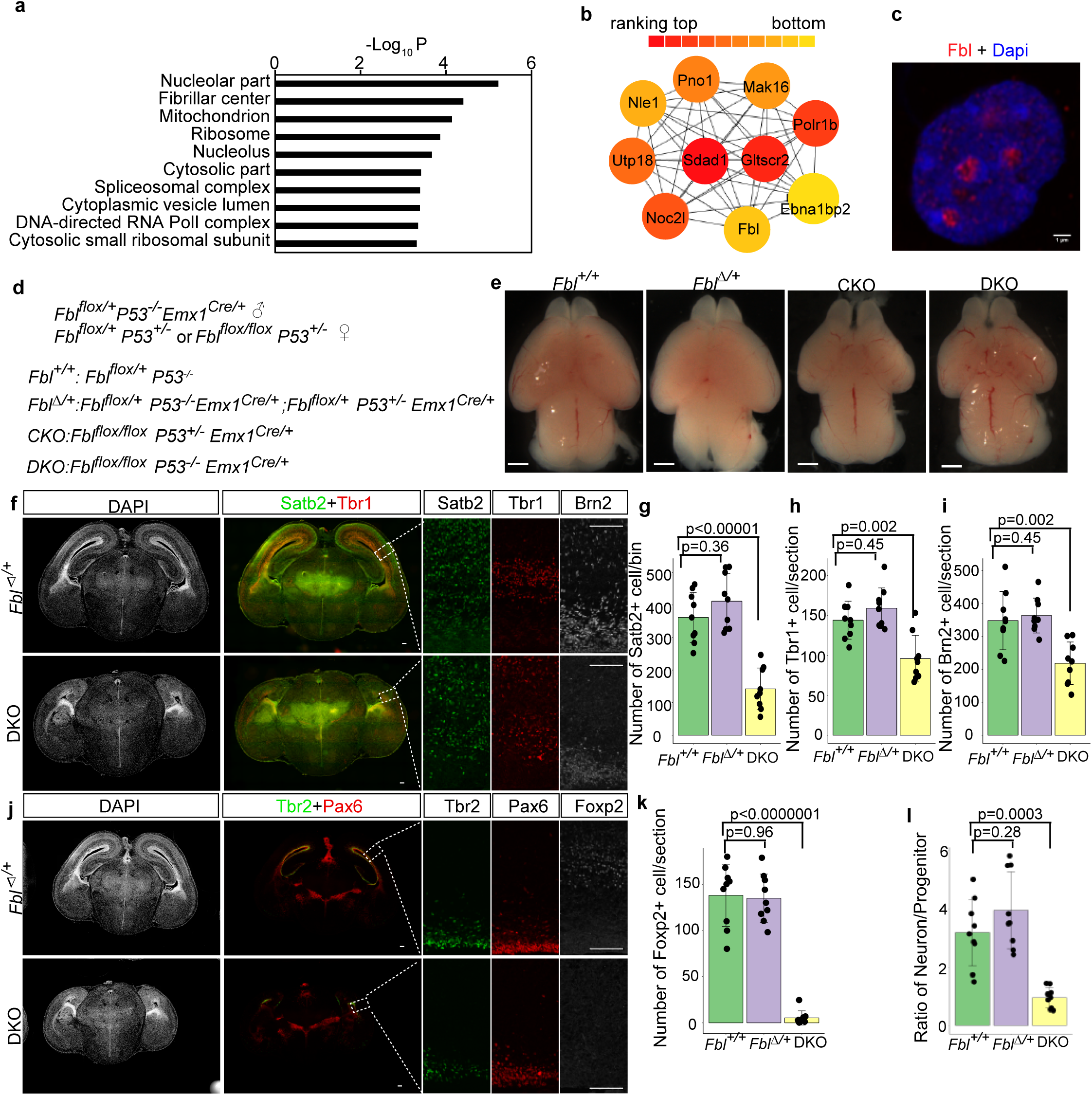
Fbl is essential for brain development. **a**, Gene Ontology (GO) analysis of the gene module (brown module) that showed higher expression in E11 than E14 NSCs. **b**, Top 10 module nodes based on protein interaction networks in the brown module. **c**, E14 NSC stained for Fbl and Dapi showing nucleolar Fbl expression (n=5). Scale bars, 1 µm. **d**, Mutant mice generation. **e**, Whole-mount brain image at E17 showing microcephaly after Fbl knockout. Scale bars, 1mm. **f,j**, E17 brain sections showing reduced number of both deep- and upper-neurons in DKO. Scale bars, 100 µm. **g-i,k**, Immunostaining-based cell number quantification (n=3 mice per genotype, n=3 sections per mouse). **l**, Ratio of Tbr1+ or Brn2+ neurons and Pax6+ or Tbr2+ progenitors (n=3 mice per genotype, n=3 sections per mouse). Data are presented as the mean±s.d. of n=9 sections (one-way ANOVA followed by Tukey’s post-hoc tests).

### Knockout of *Fbl* disrupts brain development independently of apoptosis

To examine Fbl function in NSCs, we conditionally deleted *Fbl* in the developing dorsal cortex from E9.5 by crossing with *Emx1-Cre* mice (*Fbl* CKO; Extended Data Fig. 2e). The CKO mice showed microcephaly and dramatic brain size reduction, and died around postnatal day (P) 40 (Fig. 2e; and Extended Data Fig. 2f,g). Microcephaly in *Fbl* CKO could be induced by caspase-mediated apoptosis of NSCs, since earlier work reported upregulated *Trp53* expression and subsequent apoptosis in *Fbl* knock-down mouse embryonic stem cells ^24^. Indeed, we detected high levels of cleaved caspase3 (CASP3) expression in E12.5 *Fbl* CKO brains (Extended Data Fig. 2h). To determine whether Trp53-dependent apoptosis is responsible for microcephaly, we crossed *Fbl* CKO and *Trp53-/-* mice. As *Trp53* knockout did not affect brain size, nor neuron number compared to wild type (Extended Data Fig. 3a-c), we used wild type or heterozygous mice for *Fbl* as controls (designated as *Fbl*^*+/+*^or *Fbl*^*Δ/+*^ in Fig. 2d). We obtained double-knockout mice with genotype *Fbl flox/flox, Trp53-/-, Emx1-Cre/+* (DKO) in which we confirmed the loss of Fbl by immunohistochemistry (Extended Data Fig. 2i). We also detected several rRNA sites with reduced methylations upon *Fbl* deletion, consistent with previous study showing that Fbl is a methyltransferase of rRNA^23^ (Extended Data Fig. 3d). DKO brains were smaller than control brains, although apoptosis was completely suppressed, indicating that microcephaly could not be explained by NSC apoptosis alone (Fig. 2e and Extended Data Fig. 2h).

We next tested the possibility that premature differentiation of NSCs and defective neurogenesis could cause microcephaly in *Fbl*-lacking mice by investigating DKO cortical organization at a late neurogenic stage (E17). Notably, we observed a significant decrease in the number of both deep-layer (Tbr1+ or Foxp2+) and upper-layer neurons (Satb2+) in DKO brains compared with *Fbl*^*+/+*^ or *Fbl*^*Δ/+*^ mice using immunohistochemistry (Fig. 2f-h,j,k). Moreover, the cell population expressing *Brn2*, a crucial gene for the production of upper-layer neurons ^25^, was also reduced in DKO brains (Fig. 2f,i). Additionally, we observed a significant reduction in the number of Olig2+ oligodendrocytes and of cells expressing Zbtb20, which is essential for astrogenesis ^26^ in DKO mice (Extended Data Fig. 3e-g). If premature differentiation caused a decrease in neurons, the ratio of neurons to progenitors should be increased at late neurogenesis. However, this was not the case (Fig. 2j,l), indicating that the *Fbl*-deleted mouse cortex had defective neurogenesis that was not caused by premature NSC differentiation.

### Analysis of temporal identity of *Fbl*-mutant NSCs at single cell level

To clarify the possible mechanisms leading to defective neurogenesis in DKO, we performed transcriptome analysis at the single-cell level for different genotypes along the developmental timeline (see methods for sample collection). Analysis of these transcriptome data using t-distributed stochastic neighbor embedding (t-SNE) clustered the cells according to developmental time and cell type (Extended Data Table 3a). At E14, the DKO cells were clearly separated from *Fbl*^*+/+*^or *Fbl*^*Δ/+*^ cells, but comparable to control cells at E10 or E12 (Extended Data Fig. 4a; and Extended Data Table 3b-d). We detected that differentially expressed genes (DEGs, fold change > 0.25, q-value < 0.01) between *Fbl*^*Δ/+*^ and DKO NSCs (cells in cluster 0, 1, 5, 7, 11, 12 in Extended Data Fig. 4b) increased from 41 (E10), to 74 (E12), and 760 (E14) (Extended Data Table 3b-d). These observations indicate that dramatic transcriptome changes occur in DKO NSCs compared to *Fbl*^*Δ/+*^ control NSCs after E12.

Next, we investigated the properties of DEGs at E14. As we showed previously, NSCs gradually change their identity to produce different types of neurons (Fig. 3a). Consistent with this model, PCA organizes the cells from E11 and E14 dorsal brains into two directions: a differentiation axis (PC1) and a temporal axis (PC2) (Fig. 3b). We computed the contributions of each gene to PC1 and PC2, which represent the relevance of the gene to each axis (Extended Data Table 3e). Compared with randomly selected genes, the contribution of DEGs was significantly different in both PCs, suggesting that *Fbl* deletion affects both the differentiation and the temporal axis (Fig. 3c).

**Figure 3.**
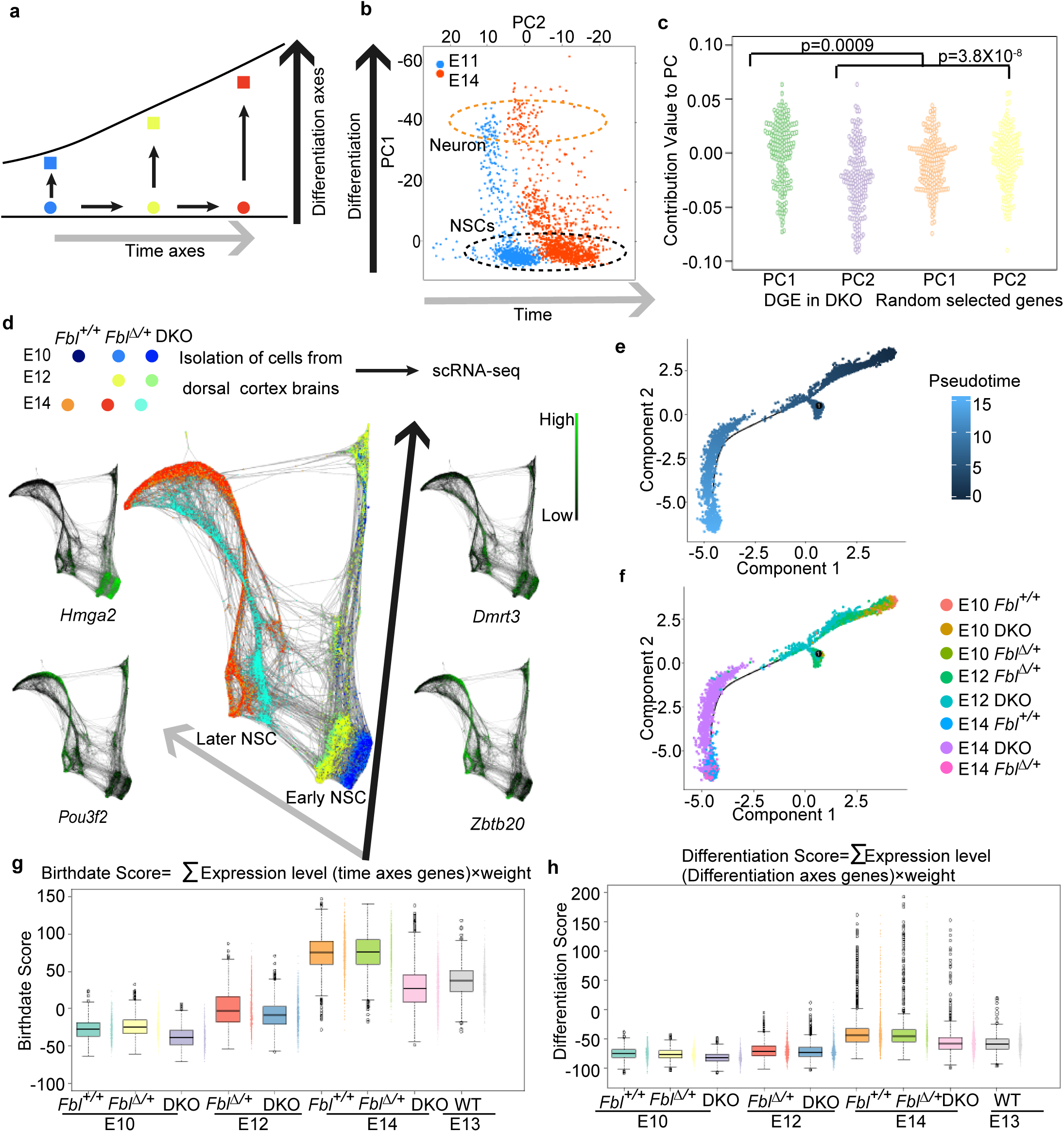
Single cell transcriptome analysis of temporal patterning in NSCs. **a**, Model of NSC temporal identity progression. **b**, Principal components analysis (PCA) of transcriptome from E11 (n=846) and E14 cells (n=1293) organizes the cells on two axes: differentiation axis (PC1) and time axis (PC2). Orange circle: neurons; black circle: NSCs. **c**, PC contribution of differentially expressed genes (DEG) between E14 DKO and Fbl^Δ/+^ NSCs showing that Fbl affects both the differentiation axis and the time axis (Kruskal-Wallis test and Dunn’s test with Bonferroni correction). **d**, SPRING graph of single cells coloured by genotype from different stages, and expression patterns of several early- and late-onset genes. E10 Fbl^+/+^ n=836, E10 Fbl^Δ/+^ n=1202, E10 DKO n=651, E12 Fbl^Δ/+^ n=2070, E12 DKO 2260, E14 Fbl^+/+^ n=3905, E14 DKO n=2879, E14 Fbl^Δ/+^ n=2592. **e,f**, Pseudo-time alignment of NSCs via Monocle. **g,h**, Scoring single-cell identity with a mathematical model. Only NCSs that identified according tSNE analysis were used in e-h (see Extend Data Fig. 4b). E10 Fbl^+/+^ n=691, E10 Fbl^Δ/+^ n=982, E10 DKO n=581, E12 Fbl^Δ/+^ n=1228, E12 DKO n=1183, E14 Fbl^+/+^ n=1046, E14 DKO n=1480, E14 Fbl^Δ/+^n=577. Data are presented as box-whiskers (left) and bee swarm plots (right).

We then asked whether Fbl promotes or suppresses the progression of NSCs along these two axes. To answer this question, we again constructed a continuous trajectory of all cells with SPRING ^16^ (Extended Data Fig. 4c). The cells were thus deposited along the differentiation and temporal axes. Consistent with the t-SNE analysis, while DKO cells from E10 and E12 could not be distinguished from controls at the same stages, E14 DKO cells were closer to E12 *Fbl*^*Δ/+*^ than to E14 *Fbl*^*+/+*^or *Fbl*^*Δ/+*^ cells, implying delayed temporal identity transition (Fig. 3d).

To further confirm our results, we introduced a simple mathematical model to estimate the developmental time and differentiation state of each NSC (cells in cluster 0, 1, 5, 7, 11, 12 in Extended Data Fig. 4b; see methods). We defined the birthdate score and the differentiation score of each cell as a weighted linear combination of specific temporal-axis and differentiation-axis genes, respectively ^12^. These scores are likely a faithful representation of each cell, as both birthdate and differentiation scores increased from E10 to E14. The birthdate scores of E14 DKO NSCs were lower than those of E14 *Fbl*^*+/+*^or *Fbl*^*Δ/+*^ NSCs, but similar to those of E13 *Fbl*^*+/+*^ NSCs (Fig. 3g). In addition, E14 DKO NSCs were less differentiated than E14 *Fbl*^*+/+*^or *Fbl*^*Δ/*+^ (Fig. 3h). Pseudotemporal ordering of NSCs from these stages also suggested a delay of temporal patterning (Fig. 3e,f). Indeed, immunohistochemistry confirmed the persistence of an early-onset gene, Dmrt3, and delayed production of later-born neurons in the E14 DKO brains (Extended Data Fig. 4d-f). These results strongly suggest that Fbl is required for the proper temporal patterning of NCSs.

### Fbl affects cell cycle progression

We next examined whether Fbl affects cell cycle progression by measuring the 5-ethynyl-2’-deoxyuridine (EdU) incorporation into NCSs. EdU pulse-labelling of S-phase cells for 1 h and immunostaining of M-phase cells with anti-phospho-histone 3 (pH3) antibody revealed significant reductions of both S-phase and M-phase cell populations in DKO at the E14 compared to control (Fig. 4a-c). To further investigate this cell cycle defect, we analyzed DNA content of NSCs using FACS after siRNA-dependent *Fbl* knockdown (Fig. 4d,e). A significant increase in NSCs at the G1/G0 phase and a reduction of S-phase NSCs were observed 2 days after *Fbl* knockdown, suggesting that Fbl impacts S-phase initiation (Fig. 4f).

**Figure 4.**
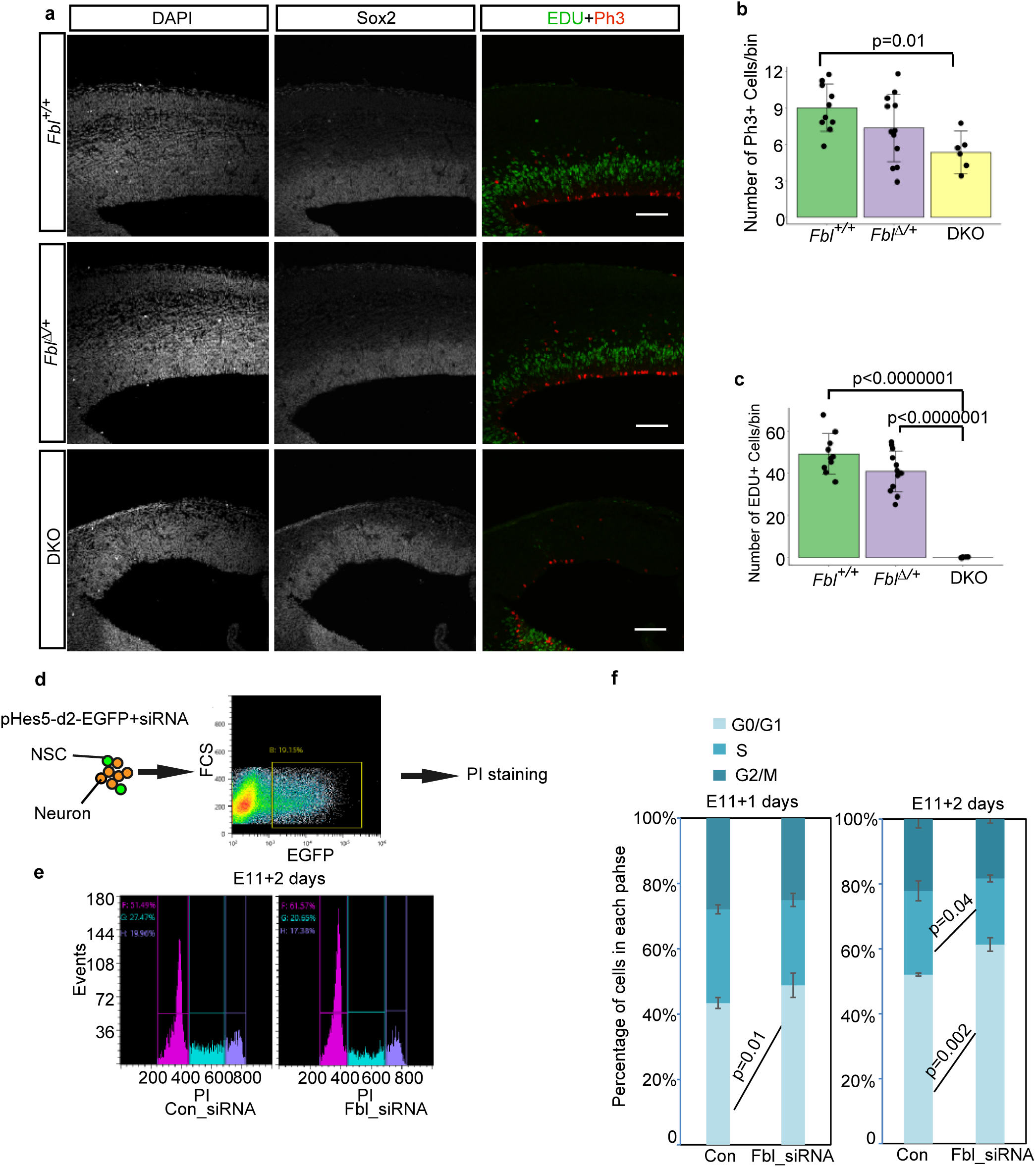
Fbl is essential for cell cycle progression. **a**, Representative image of E14 brain sections stained for Sox2, Edu, and Ph3. Edu was injected 1 h before sampling. Scale bar: 100 µm. **b, c**, Cell number quantification on sections based on immunostaining with the indicated markers (n=5, 6, and 3 mice for *Fbl*^*+/+*^, *Fbl*^*Δ/+*^, and DKO, respectively; n=2 sections per mouse; one-way ANOVA followed by Tukey’s post-hoc tests; data are presented as mean±s.d. of counted sections). **d**, Schematics of the experimental design to investigate NSC cell cycle progression after treatment with control or *Fbl* siRNA. *pHes5*-d2-EGFP was used to label NSCs. **e**, Cell cycle analysis of NSCs after *Fbl* knockdown. **f**, Proportion of G1/G0, S, and G2/M phase change after *Fbl* knockdown for 1 day (left) or 2 days (right) (Student’s t-test; data are presented as mean±s.d.)

### Effect of Fbl on temporal identity transition is cell-autonomous

The transition in NSC identity that promotes the shift to late-born neurons from early-born neurons requires feedback interaction between NSCs and early-born neurons ^27^. However, if NSCs can receive the feedback from neighboring early-born neurons, the temporal pattern of these NSCs proceeds normally, even though their cell cycle is artificially arrested ^10^. In the case of DKO, defects in temporal patterns might come from a compromised feedback from early-born neurons. We then examined how *Fbl*-deleted cells are affected in the presence of early-born neurons in two different conditions: sparse culture of *Fbl*-deleted cells with surrounding normal cells and *in vivo Fbl* deletion in a sparse population using CRISPR/Cas9. In both cases, *Fbl*-deleted cells produced less late-born neurons than control cells (Extended Data Fig. 5). These results indicate that defective Fbl cell-autonomously compromises temporal identity progression even in the presence of normal neighboring early-born neurons, where their feedback signal can proceed temporal pattern of NCS. As cell-cycle arrested NCSs can proceed with their temporal pattern due to feedback from neighboring neurons ^10^, a cell-autonomous effect other than cell cycle defects appear to compromise the temporal pattern in Fbl-deficient NSCs.

### Fbl is essential for translation but not transcription of epigenetic modifiers

We investigated how Fbl affects temporal identity progression in NSCs. Considering that Fbl is an rRNA methyltransferase, we tested whether *Fbl* knockout affects global protein synthesis by quantification of O-propargyl-puromycin (OPP) incorporation into nascent proteins (see methods). We found decreased levels of newly synthesized proteins in DKO NSCs (Extended Data Fig. 6a). To investigate whether Fbl affects the translation of selected mRNAs, we performed ribosome profiling ^28^ and RNA-seq using *Fbl*^*Δ/+*^ and DKO brains (Fig. 5a).

**Figure 5.**
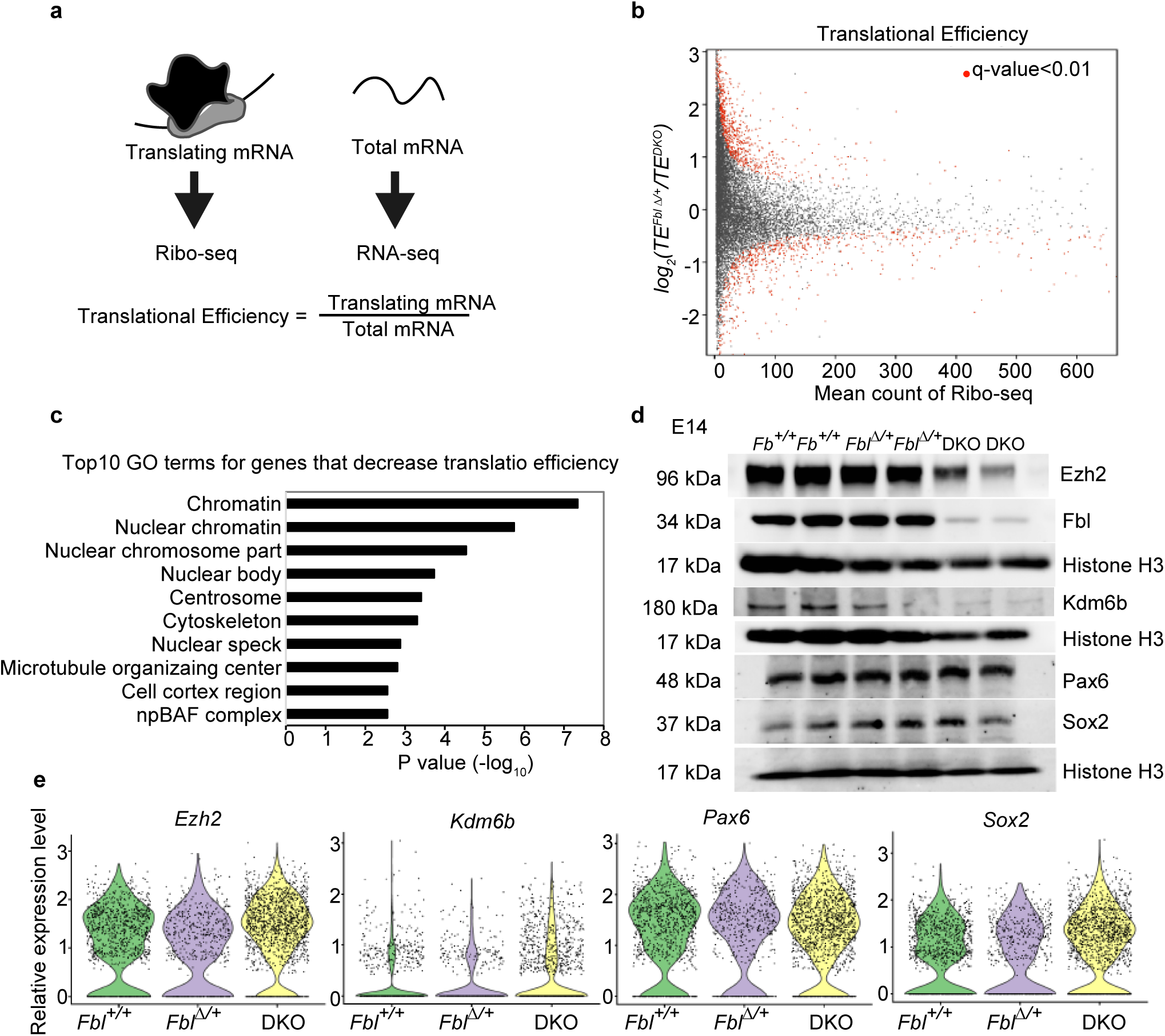
Fbl selectively regulates the translation of genes involved in H3K27me3 modification. **a**, Schematics of the experimental design for evaluation of translational efficiency (TE). **b**, Mean-count and mean-difference plots comparing observed and expected variance in TE. Genes with q-value <0.01 are shown in red. **c**, Top 10 GO terms of transcripts showing reduced TE. **d,e**, Western blotting (d) and single-cell RNA analysis (e) of the indicated genes, showing reduced protein levels, but not mRNA levels, of Ezh2 and Kdm6b in DKO brains at E14. Notice that Sox2 and Pax6 did not show changes in neither protein nor mRNA levels.

Translational efficiency (TE) can be calculated by comparing the levels of translating mRNA (Ribo-seq) and total mRNA (RNA-seq). After analyzing the qualities of data according to several criteria (Extended Data Fig. 6b-d), we detected 299 and 541 genes with an increased and decreased TE (q-value < 0.01) in DKO brains, respectively (Fig. 5b,c; Extended Data Fig. 6e and Extended Data Table 4). Given that *Fbl* deletion reduced global levels of newly synthesized protein, we considered that mRNAs with a lower TE in DKO could be directly regulated by Fbl. Indeed, Gene Ontology (GO) analysis showed that genes involved in cell cycle progression, such as *Cdk1* and *Cdk6*, were listed in the ‘centrosome’ cluster as genes with a decreased TE in DKO (Fig. 5c, Extended Data Fig. 6f,g; and Extended Data Table 4), and hence cell cycle defects might be ascribed to a lower TE of those genes (Fig. 4). Strikingly, chromatin-related genes were highly enriched among those with a decreased TE in DKO, consistent with the roles of epigenetic modifications in temporal patterning ^29^ (Fig. 5c and Extended Data Table 4). Among these highly enriched genes were *Ezh2* and *Kdm6b*, which encode a methyltransferase and a demethylase, respectively, of H3K27me3 (Extended Data Fig. 6h,i). We found that *Kdm6b* and *Ezh2* protein levels were reduced without a concurrent decrease in mRNA levels, while protein and mRNA levels of NSC markers Pax6 and Sox2 were unaffected (Fig. 5d,e). This difference was not ascribed to protein stability (Extended Data Fig. 6j), thus we deduced Fbl to be selectively targeting protein translation.

### *Fbl* affects temporal patterning though H3K27m3 modification

We investigated histone modification changes upon *Fbl* deletion in DKO and control samples (including NSCs and progenies; Fig. 6a) and demonstrated alterations of H3K27me3 and H3K4me3 modifications at 669 and 0 sites (q-value < 0.05), respectively, indicating significant defects in H3K27me3 marks (Fig. 6b,c; and Extended Data Table 5a). Moreover, the abundance of H3K27me3 peaks on early-onset genes did not show a clear difference between control and DKO samples (Fig. 6d). In contrast, compared with control samples, the abundance of H3K27me3 peaks on early-onset genes (especially on transcription start sites) was higher in DKO samples, indicating the expression of these genes was repressed by H3K27me3 (Fig. 6e). Indeed, the expression of these genes were not upregulated in NSCs of E14 DKO (Extended Data Fig. 4d). To investigate whether H3K27me3 defects in DKO can represent the temporal identity delay, we plotted these H3K27me3 samples together with previous samples (in Fig. 1m) using PCA. E14 DKO and control samples were deviated from E14 NSCs in the PCA plot due to contamination of differentiated progenitors. However, as for the temporal direction, E14 DKO samples were not clustered with cells from E14 *Fbl*^*+/+*^ nor *Fbl*^*Δ/+*^, but rather were close to E12 NSCs, reflecting a delay of temporal identity progression (Fig. 6f).

**Figure 6.**
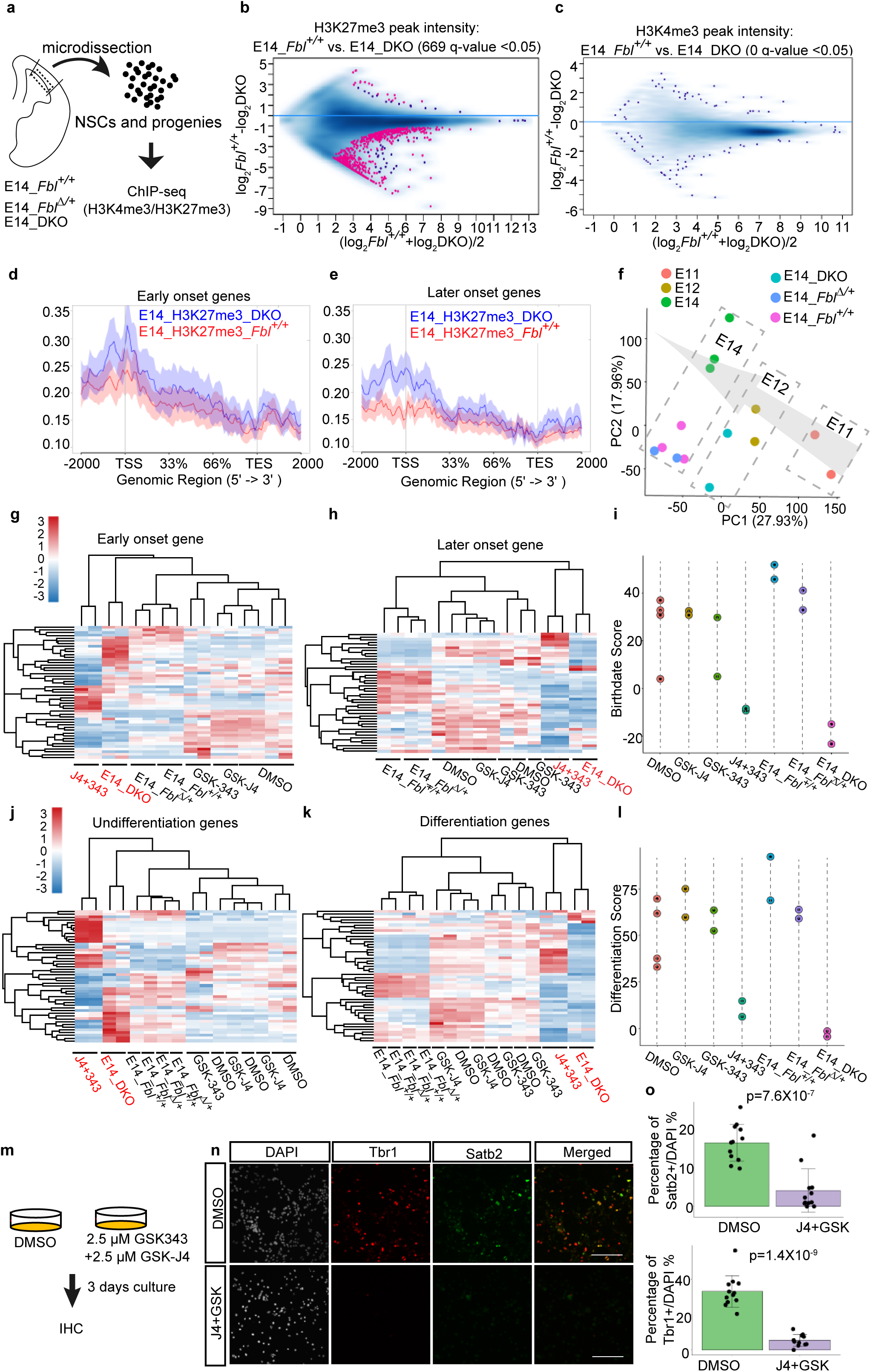
Fbl regulates H3K27me3 pattern in NSCs. **a**, Experiment design for ChIP-seq using different genotypes. **b,c**, Intensity comparison of H3K27me3 (b) and H3K4me3 (c) peaks between E14 control and DKO samples. **d,e**, Read-density profiling of H3K27m3 at early-onset (d) and late-onset genes (e) in E14 control and DKO samples. **f**, Principal component analysis (PCA) of H3K27me3 peaks, showing DKO samples were located with E12 samples. Arrow indicates temporal axes. **g-k**, Heatmap showing gene expression change after inhibitors treatment and knockout of Fbl. Core genes involvoed in birthdate and differentiation were used. In all cases, simultaneous inhibition of H3K27me3 methyltransferase and demethylase showed similar gene expression pattern with Fbl DKO. **i,l**, Birthdate (i) and differentiation (l) scoring after simultaneous inhibition of H3K27me3 methyltransferase and demethylase, showing delayed temporal progression and impeded neurogenesis. E14 samples with the indicated genotypes were used for comparison. **m**, Schematics of the experimental design to test the effects of inhibitors on NSCs. **n**, Representative image of cultured cells stained for Tbr1 and Satb2 after treatment with inhibitors. Scale bar: 100 µm. **o**, Quantification of cell number on sections based on immunostaining with the indicated markers (n=3 independent experiments). For each experiment, four regions were randomly selected and counted (Student’ s t-test; data are presented as mean±s.d. of n=12 counted images).

Fbl control of both *Ezh2* and *Kdm6b* led us to questioning the role of modification turnover during temporal patterning. To this end, we inhibited both methyltransferase and demethylase using the specific inhibitors GSK-343 and GSK-J4, respectively, investigated gene expression changes involved in birthdate and differentiation, and calculated the birthdate and differentiation scores after RNA-seq. Inhibition of H3K27me3 methyltransferase or demethylase alone affected the expression of 210 and 409 genes, respectively (q-value<0.05); however, it did not dramatically affected birthdate and differentiation scores (Fig. 6g-l). In contrast, the simultaneous suppression of methyltransferase and demethylase affected the expression of 10056 genes (q-value<0.05) and reduced both scores to levels similar to that of DKO brains, indicating delayed temporal fate progression (Fig. 6g-o; Extended Data Table 5b-e). Therefore, we concluded that both writing and erasing of H3K27me3 are essential for temporal pattering, and that Fbl facilitates these processes by controlling the translation of key enzymes.

### Fbl selectively enhances the translation of targets genes though 5′UTRs in a cap-dependent manner

Finally, we investigated how Fbl promotes the translation of target mRNAs. As the 5′untranslated region (UTR) plays a critical role in translational regulation, we analyzed the 5′UTRs of Fbl target mRNAs showing downregulated TE after *Fbl* deletion. Cap-independent translational initiation by RNA regulons such as internal ribosome entry site (IRES) elements can confer translational selectivity to specific mRNAs. IRES elements are characterized by the presence of poly (U) motif or a highly organized secondary structure ^30^. The minimum free energy (MFE) of the 5′UTRs of Fbl target mRNAs was significantly lower than that of randomly selected mRNAs, suggesting that they were more structured (Fig. 7a). Moreover, a poly (U) motif was highly enriched in the 5′UTRs of these target mRNAs (31 genes; p-value=10^−56^, Fig. 7b). Therefore, it is highly likely that the translation of Fbl target mRNAs is driven by a cap-independent mechanism in NSCs. To test this hypothesis, we constructed bicistronic reporters, in which cap-independent translational activity can be measured by the ratio between BFP and GFP, which are translated in a cap-dependent and a cap-independent manner, respectively (Fig. 7c). As a positive control, we used the 5′UTR of *Cdkn1b*, which is a known cellular IRES ^31^. We detected GFP signals by FACS when the 5′UTRs of *Cdkn1b, Kdm6b* and *Ezh2* were used. Knockdown of *Fbl* reduced these cap-independent translational activities, while BFP levels were not changed (Fig. 7d and Extended Data Fig. 7a-d). In contrast, only background level of GFP signals was observed when the 5′UTR of *Pax6* was used, and the GFP signal did not depend on *Fbl* dosage (Extended Data Fig. 7a).

**Figure 7.**
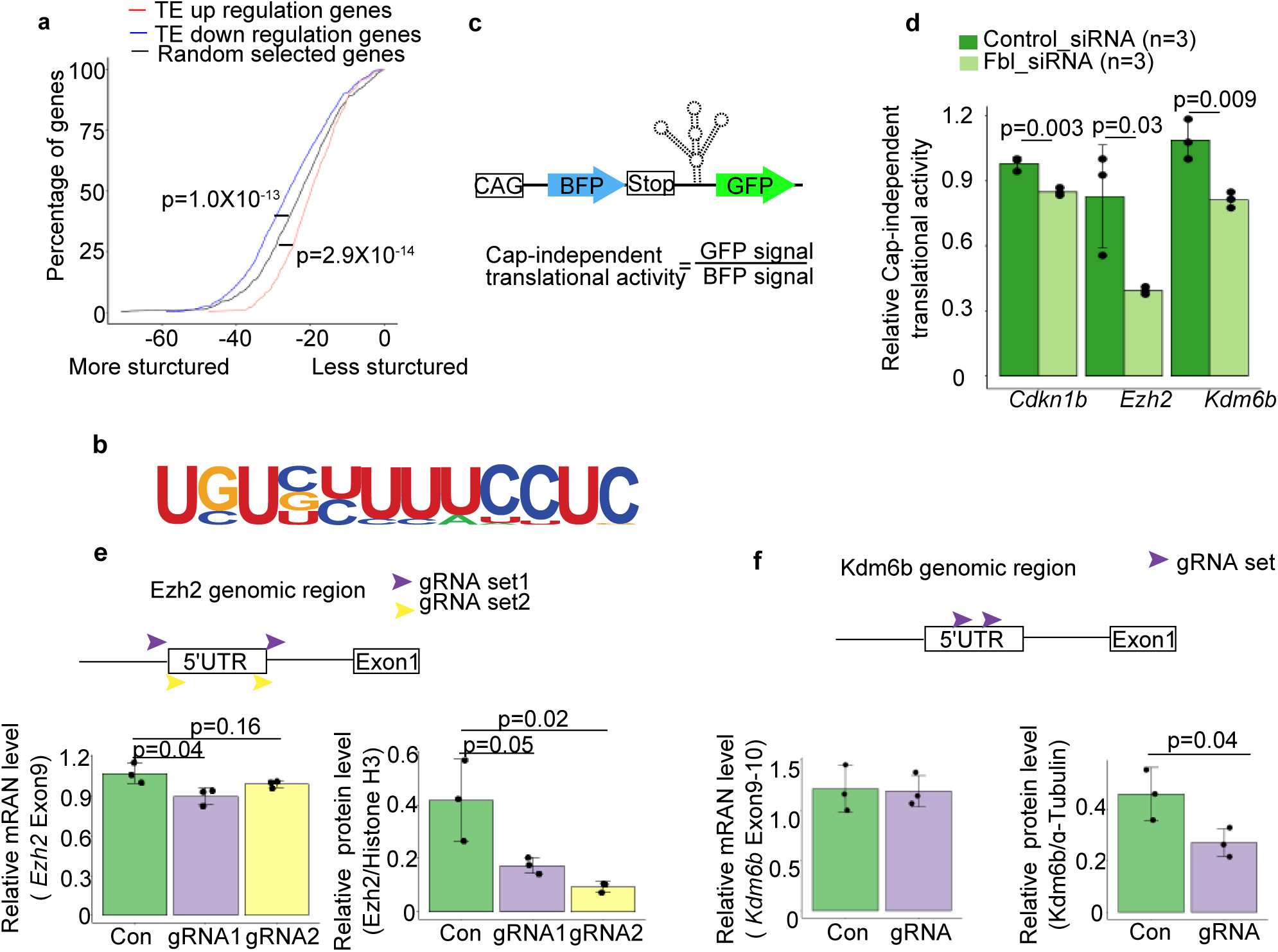
Fbl regulates translation through the 5′UTR in a cap-independent manner. **a**, 5′UTR minimum free energy (MFE) cumulative distribution of mRNAs showing changes in translational efficiency (TE) after Fbl knockout. Randomly selected mRNAs are shown as controls (Wilcoxon signed rank test). **b**, A poly(U) motif enriched in the 5′UTRs of mRNAs with downregulated TE. **c**, Experimental assessment of cap-independent translational initiation. **d**, Relative cap-independent translational activity in control and Fbl knockdown cells (n=3, Student t-test; data are presented as mean±s.d.) **e,f**, Changes in mRNA (left) and protein levels (right) after Ezh2 (e) and Kdm6b (f) 5′UTR knockout (n=3; one-way ANOVA followed by Tukey’s tests (e) and Student t-test (f); data are presented as mean±s.d.)

Furthermore, to uncover the role of *Ezh2* and *Kdm6b* 5′UTRs *in vivo*, we disrupted their 5′UTRs in NSCs using CRISPR/Cas9 (Extended Data Fig. 7g,h), resulting in significantly reduced protein levels, with mRNA levels either slightly or not affected (Fig. 7e,f; Extended Data Fig. 7e,f). These results strongly suggest that Fbl enhances translation of these genes through 5′UTR. However, we could not confirm this 5′ UTR mechanism as identical with well-known viral IRES, a cap-independent 5′UTR translation initiation, because *Ezh2* and *Kdm6b* could still be translated with a low efficiency in the absence of *Fbl* or 5′UTR (Fig. 5d; and Extended Data Fig. 7e,f).

## Discussion

In this study, we showed that Fbl drives the H3K27me3-dependent developmental clock independent of cell cycle. As a simple model to measure developmental time, cell cycle number can perfectly predict the initiation of transcription from zygotic genome during early development of *Xenopus* (12 cycle) and *Drosophila* (10 cycle) ^2 3^. However, many studies show that the developmental clock can work independent of cell cycle progression. As described before, p27/kip1 accumulation during proliferation is important for differentiation of oligodendrocyte precursors; however, slow down cell cycle progression of oligodendrocyte precursors by culture them in 33 °C rather than 37 °C does not affect differentiation process ^32^. Moreover, during sequential expression of four temporal fate genes (*hunchback, Krüppel, pdm* and *castor*) in *Drosophila* neuroblasts, the *hunchback* to *Krüppel* transition required cytokinesis, but the sequential expression of *Krüppel, pdm*, and *castor* is observed in the G2-arrest neuroblasts, indicating that their progression is independent of cell cycle ^33^.

We observed both H3K4me3 and H3K27me3 peaks in the genomic region of early onset genes (Fig.1g,i). This bivalent state is considered a mechanism for maintaining a potential for genetic activation ^34^. This bivalent state of early-onset genes may also explain the plasticity of late NSCs, in which reprogramming of temporal identity has been reported after transplantation into young brains ^35^. Unlike H3K4me3, we observed a dramatic gain and loss of H3K27me3 modification in NSCs between E11 to E14. Since H3K27me3 is highly associated with topologically associated domains and chromatin subcompartments ^36 37^, this result strongly suggests that global chromatin organization, rather than repression of specific genes, is essential for temporal patterning of NSCs. Consistently, the simultaneous inhibition of methyltransferase and demethylase of H3K27me3, which mimics the *Fbl* loss-of-function phenotype, yielded much more DEGs (10056) than the linear addition of two inhibitors working separately (210 and 409, respectively; Extended Data Table 5).

Recently, 2′-*O*-methylation sites were found in mRNAs ^38,39^ where they inhibit translation elongation by slow tRNA decoding ^40^ rather than facilitating translation as found in this study. Thus, it is more likely that rRNA, rather than mRNA, modifications mediate the effect of Fbl on translation during brain temporal patterning. We hypothesized that rRNA modification by Fbl facilitates ribosomes to recognize or bind 5′UTR of target genes, thereby enhancing their translation (Fig. 8). In addition, it is likely that Fbl affects translation via the structure of 5′UTR of target mRNAs, which restricts the range of translational regulation, eventually generating the specificity of Fbl targets. Moreover, some ribosomal protein isoforms also have differential preferences for translation of specific mRNAs ^41^. Indeed, scRNA data indicates that a subset of ribosomal proteins showed higher expression in early compared to late NSCs (Fig. 2a) ^42^. Therefore, ribosomal proteins and rRNA modification likely work coordinately to ensure the precise control of translation.

**Figure 8.**
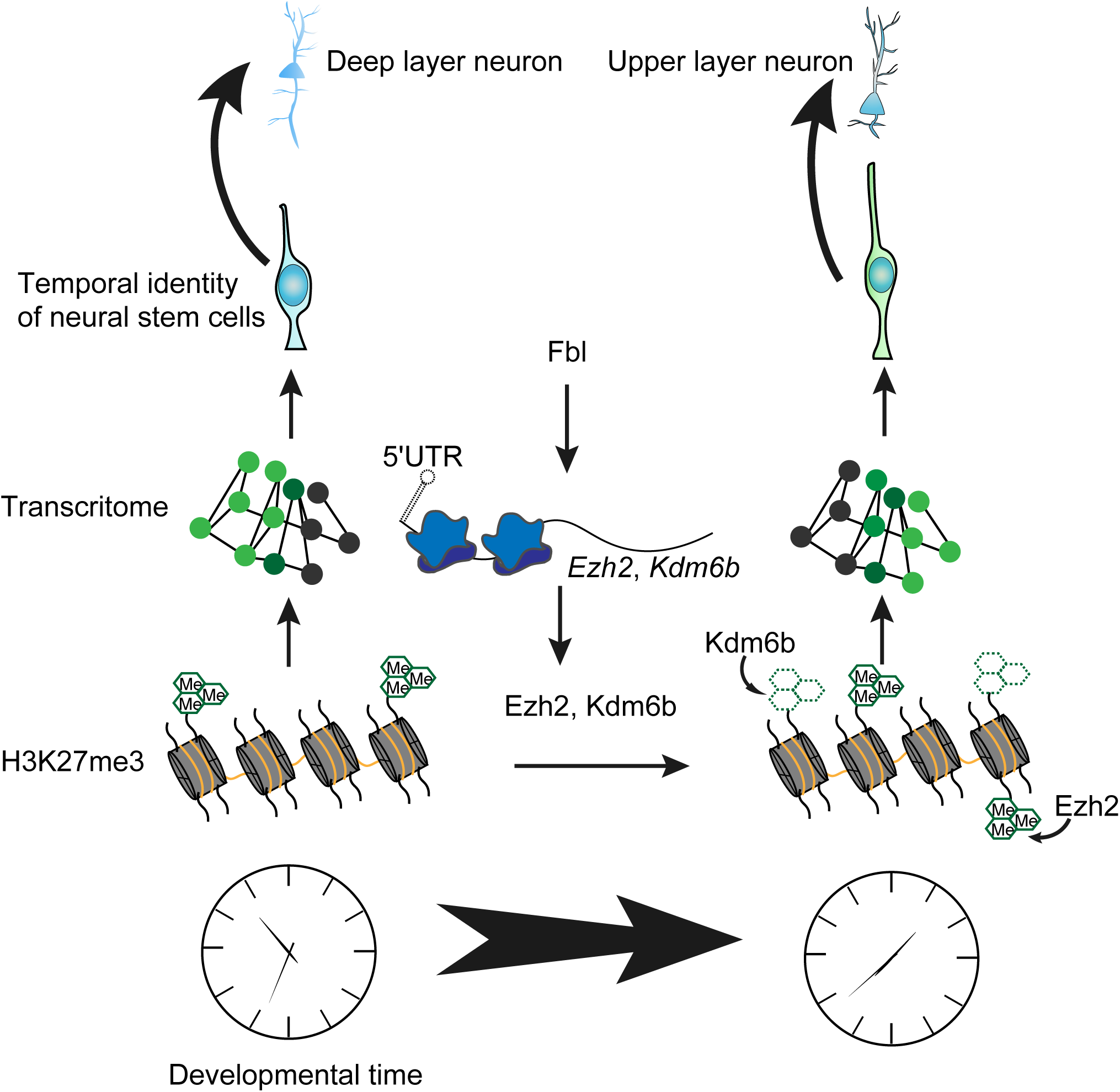
Fbl drives developmental clock of NSCs. Fbl selectively enhances the translation of *Ezh2* and *Kdm6b* through their the 5′UTR in a cap-independent manner. Ezh2 and Kdm6b change H3K27me3 pattern in NSCs. H3K27me3 patterning further affects gene expression change and regulates the temporal fate of NSCs.

Why does epigenome-mediated temporal patterning utilize translational control by Fbl as an upstream mechanism to advance the developmental clock, in addition to transcriptional control of epigenetic factors? We speculate that selective translational promotion through 5′UTR is simple and efficient in ensuring the production of specific protein groups; the transcriptional upregulation of a group of genes often needs the cooperation of many epigenetic and transcription factors. Translational control of epigenetic modifiers by Fbl adds another level of complexity to gene expression, and thus, greatly widens the range of the epigenetic landscape, notably impacting diverse developmental processes and diseases.

## Materials and Methods

### Animals

All animal procedures were performed in accordance with the guidelines for animal experiments at RIKEN Center for Biosystems Dynamics Research.

### Mice

To produce conditional knockout mice of *Fbl* (Accession No. CDB0137E: http://www2.clst.riken.jp/arg/mutant%20mice%20list.html), *loxP* sequences were introduced on both sides of *Fbl* locus by genome editing technology using CRISPR/CAS9 system in mouse zygotes. crRNA(CRISPR RNA), tracrRNA(trans-activating crRNA) and donor single-stranded oligodeoxynucleotides (ssODNs) consisting of a *loxP* site, an EcoRV recognition site and homology arms were chemically synthesized (Fasmac): 5′-crRNA (5′-AGCUUGUCUCAGGUUUAACCGUUUUAGAGCUAUGCUGUUUUG −3′), 3′-crRNA (5′-UCAAGGGCGCAUGCGUCUCGGUUUUAGAGCUAUGCUGUUUUG-3′) and tracrRNA (5’-AAACAGCAUAGCAAGUUAAAAUAAGGCUAGUCCGUUAUCAACUUGAAA AAGUGGCACCGAGUCGGUGCU-3’), 5′-*loxP* ssODN (5′-GTCCTCAGCACACAGCTTGTCTCAGGTTTAGATATCATAACTTCGTATAGCATACATTATACGAAGTTATACCTGGTTCCACATCACACCTGCCGCTGTT-3′) and 3′-*loxP* ssODN (5′-CACACAAAGTTGATCAAGGGCGCATGCGTCATAACTTCGTATAATGTATGCTATACGAAGTTATGATATCTCGAGGCCACTTAGCAATAGGCACCAGACA-3′). The mixture of ssODNs, crRNAs, tracrRNA and Cas9 protein was injected into pronuclei of C57BL/6N zygotes by microinjection, and the injected zygotes were transferred into the oviducts of pseudopregnant ICR female mice. The resultant offspring were genotyped by genetic PCR with combination of following primers: 5′-*loxP* site: forward: 5′-CTCTTCTAGGACACTCCATCCCTTATCAAG-3′; reverse: 5′-AGTACTAGTTGTGAAGGTATGAGAGGGGTC-3′; (wild type: 489 bp and 5′*loxP*: 529 bp) 3′-*loxP* site: forward: 5′-GAAGAAGATGCAGCAGGAGAACATGAAGCC-3′; reverse: 5′-CAACCAGCAAAATGGCGACCACAACAAACC-3′ (wild type: 575 bp and 3′*loxP*: 615 bp). The insertions of *loxP* site were confirmed by the EcoRV digestion (5′-*loxP*: 292 bp and 237 bp, and 3′-*loxP*: 360 bp and 255bp) and sequencing. The germline transmission of floxed allele in which 5′-and 3′-*loxP* sites were in the same allele was confirmed by crossing with C57BL/6. Production of *Trp53* mutant mice is described before ^43^.

*Fbl* and *Trp53* mutant mice was maintained in C57BL/6 background. The reporter mouse line: *pHes1*–*d2-EGFP* (a gift from R. Kageyama ^17 44^) was maintained in ICR background. Wild type mice used for inhibitors treatment and cap-independent translational activity tests were maintained in ICR background.

### Weighted Gene Coexpression Network Analysis (WGCNA)

WGCNA was performed using microarray data of E11 (n=24) and E14 (n=31) single cells ^10^. We used top 10,000 genes after ranking of all genes by their expression level. Soft power parameter was set at three and dpSplt parameter was set as 0. Genes in the brown module were used to analyze protein interaction (https://string-db.org) and top10 nodes were identified by cytoHubba in Cytoscape ^45^. Gene enrichment analysis was performed using Enrichr ^46^.

### Immunohistology and confocal imaging

Brains were fixed in 1% paraformaldehyde (PFA) overnight, treated by 25% sucrose for cryoprotection, and then embedded in OCT compound (Tissue-Tek; Sakura). Sections (12 μm) were made using a cryostat (CM3050S Leica Microsystems). For immunostaining, sections were blocked with 3% skim milk powder in PBST (0.1% Tween20 in PBS) for 1 h at room temperature (RT), followed by incubation with primary antibody in the optimized concentration at 4 °C. According to primary antibodies, sections were then incubated with secondary antibodies with labelled fluorescent probes (1:400) (Alexa Flour 488, cy3, or 647; Jackson ImmunoResearch) for 90 min at RT. DAPI was used for nuclei detection. A scanning confocal microscope (Olympus FV1000 or Zeiss LSM 880 with Airyscan) was used for observation. For Tbr2/EOMES staining, the antibody (rat monoclonal, clone Dan11mag; eBioscience at Thermo Fisher) has been conjugated with eFluor660. The primary antibodies were: Fibrillarin (rabbit polyclonal, ab5821, abcam), Cleaved Caspase-3 (Asp175)(rabbit polyclonal, 9661S, Cell Signaling Technology), Satb2 (mouse monoclonal, ab51502, abcam), Tbr1 (rabbit polyclonal, ab31940, abcam), Olig2 (goat polyclonal, AF2418, R&D System), Pax6 (rabbit polyclonal, PRB-278P, Covance), Sox2 (goat polyclonal, sc-17320, Santa Cruz), GFP (chick polyclonal, GFP-1020, aves), Phospho-Histone H3 (Ser10) (rabbit polyclonal, 06-570, Millipore), Brn-2 (goat polyclonal, sc-6029, Santa Cruz), FoxP2 (goat polyclonal, sc-21069,, Santa Cruz), Zbtb20 (Rabbit polyclonal, HPA016815, Sigma) and Dmrt3 (rabbit polyclonal, a gift from D. Konno) ^47^.

### Western blot analysis

Pierce® IP lysis buffer (Thermofisher Scientific) was added into the collected dorsal cortices or the FACS-sorted cells. After sonication (TOMY HandySonic, 10s at level 4), lysate was centrifuged at 13,000 g for 10 min. The supernatant was mixed with 1 µl of protease inhibitor cocktail (nacalai tesque). After mixing with same volume of sampling buffer laemmli (Sigma), the supernatant was boiled at 98 °C for 5 min, applied to SDS page gel (SuperSep, WAKO) and transferred onto a 0.2 µM nitrocellulose blotting membrane (GE Healthcare). After incubation with blocking buffer (5% milk in tris-buffered saline with 0.1% Tween20) at RT for 1 h, the membrane was incubated with primary antibody overnight at 4 °C, followed by the incubation with anti-rabbit or anti-mouse IgG antibody conjugated to horseradish peroxidase (NA934V or NA931V, GE Healthcare). After reactivation with Chemi-Lumi One Ultra (nacalai tesque), images were obtained by LAS3000 mini imaging system (Fujifilm). The intensity of the bands was calculated by ImageJ (1.52d). The primary antibodies were: Ezh2 (mouse monoclonal, 5246S, Cell Signaling Technology), Pax6 (rabbit polyclonal, PRB-278P, Covance), Histone H3 (rabbit monoclonal, 4499, Cell Signaling Technology), Kdm6b (rabbit polyclonal, NBP1-06640, Novus Biologicals), Sox2 (rabbit polyclonal, ab75179, Abcam), Fibrillarin (rabbit polyclonal, ab5821, Abcam) and α-tubulin (mouse monoclonal, clone DM1A, T9026, Sigma-Aldrich).

### Plasmid, stealth siRNA and gRNA

Stealth siRNA for *Fbl* and control was designed with BLOCK-iT™ RNAi Designer (https://rnaidesigner.thermofisher.com/rnaiexpress/) as following: *Fbl* siRNA: 5′-CCGCAUCGUCAUGAAGGUGUCUUUA-3′ and control siRNA: 5′-CCGGCUACUAGUGGAUGUUCCAUUA-3′.

To construct reporter of cap-independent translational activity, sequence of mtagBFP, 5′UTR of test genes and EGFP were inserted into pCAG-FLAG-N1 plasmid ^48^ using In-Fusion HD Cloning Kit (TAKARA).

The *pHes5-d2-EGFP* plasmid used in cell cycle analysis was a gift from R. Kageyama ^17 44^. The target region of gRNA for knockout of *Fbl* and *Trp53* was showed below: *Fbl* gRNA1: 5′-CCACCATGCGGCATGCTGGAATT-3′; *Fbl* gRNA2: 5′-CCTCGAGACGCATGCGCCCTTGA-3′; *Trp53* gRNA1: 5′-CCTCGCATAAGTTTCCTGAAATA-3′; *Trp53* gRNA2: 5′-CAGCAGGTGTGCCGAACAGGTGG-3′;

The target region of gRNA for knockout of 5′UTR of *Ezh2* and *Kdm6b* was showed below: *Ezh2* gRNA set1_1: 5′-GGGTTGCTGCGTTTGGCGCTCGG-3′; set1_2: 5′-CCGTCGGCCGCCGGTGGTCGGCA-3′; *Ezh2* gRNA set2_1: 5′-CCGGTCGCGTCCGACACCCAGTG-3′; set2_2: 5′-GAGAGGCGCGGGCTGGCGCGCGG-3′.

*Kdm6b* gRNA set1_1:5′-CCCTCAGGTCGGCTCGTGAATGG −3′ set1_2: 5′-CCCACTTGCGCGATTCTAGGGGC-3′. Primers for production of gRNA were designed following ^49^. Designed primers were self-amplified by PCR and PCR fragments were inserted into AflII cut gRNA vectors modified from Church Lab ^50^.

All plasmids were purified with endotoxin-free NucleoBond Xtra Midi EF kit (Macherey– Nagel).

### EDU staining

Pregnant mice were injected intraperitoneally with EdU (Invitrogen) at 12.5 mg/kg. EdU staining was performed with Click-iT EdU Imaging kit (Invitrogen).

### Single cell isolation and library construction

To isolated *Hes1* positive neural stem cells (NSCs) at E14, dorsal cortices were dissected from *Hes1-d2-EGFP*^*Tg/+*^ mice. Cortices were dissociated with 0.05% trypsin with Hanks’Balanced Salt Solution (HBSS) (–) at 37 °C for 10 min. After centrifugation at 1000 g for 5 min, cells were resuspended with 0.375% BSA/HBSS(-) by gentle pipetting 15 to 20 times. Resuspended cells were filtered with 35 µm filter (Falcon) and sorted into sorting buffer (20 ng/ml human basic FGF (Peprotech), 1XB27 RA-(Gibco), in Dulbecco’s Modified Eagle Medium (DMEM) F12+GlutaMax (Gibco)) by a cell sorter (SH800, SONY) equipped with 130 µm sorting chips (SONY, LE-C3113). After sorting, cell number was counted by Countess or Countess II (Invitrogen). For cells from wild type and *Fbl* mutant mice, cells were collected similarly except that sorting was not performed.

Collected cells were immediately load into the 10X-Genomics Chromium (10X Genomics, Pleasanton, CA). Libraries for single cell cDNA were prepared using Chromium 3′ v2 platform as the manufacturer′s protocol.

### Bioinformatics analysis of single cell data

Sequenced data was mapped to mm10 and cell number and raw count for each gene were reported by cellranger 2.0.2 (10X GENOMICS). All data were further analyzed by a bioinformatics pipeline Seurat (2.3.4) ^51^. Briefly, we first created a SeuratObject, in which genes that were expressed by less than 3 cells and cells that expressed less than 200 genes were removed. Number of unique molecular index (UMI) were automatically counted by Seurat. We calculated percentage of UMI mapping to mitochondrial genes and used these values to further filter cells (<15%). Data were then normalized ((normalization.method = “LogNormalize”, scale.factor = 100000) and scaled (vars.to.repress=c(“UMI”,”percent.mito”)). Then, principal component (PC) analysis were performed and top 10 PCs were used for t-Distributed Stochastic Neighbor Embedding (t-SNE) dimensional reduction. TSNEPlot and FindAllMarkers was used to visualize clusters and to investigate the cell type of each cluster, respectively.

### Pseudotemporal ordering

To order NSCs pseudotemporally, monocle (2.10.1) package was used ^52^. NSCs were extracted based on t-SNE clustering (Cluster 0, 1, 5, 7, 11, 12 in Extended Fig. 4b). NSCs were manually clustered into early NSCs and later NSCs according to the expression level of *Hmga2* (>2) and *Dbi* (>15). Then, these cells were ordered by DDRTree method according 200 differential expression genes between these two clusters.

### Calculation of birthdate and differentiation score

To evaluate temporal identity and differentiation state of each cell, we used core genes to calculate birthdate and differentiation score as previous reported ^12^. Briefly, authors used ordinal regression models to predict birthdate and differentiation state of each cell. The best 100 genes based on the linear weight of the models was selected for prediction. We used 95 genes (5 of them could not be detected in our system including Rp23-379c24.2, Leprel1, Rp23-14p23.9, Mir99ahg and Yam1) and 100 genes for calculation of birthdate and differentiation score by following formula:

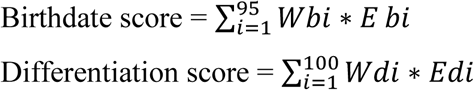

*Wb*_*i*_ and *Eb*_*i*_ indicates weight of each temporal-related gene and its expression level in each cell, respectively. *Wd*_*i*_ and *Ed*_*i*_ indicates weight of each differentiation-related gene and its expression level in each cell, respectively.

### Ribosome profiling

Ribosome profiling protocol was modified from previous study ^53^. Briefly, E14 dorsal cortices were dissected and removed into a 1.5 ml tube. Then, 200 µL and 100 µL (in the case of *Fbl*^*Δ/+*^ and DKO cortices, respectively) ice-cold lysis buffer (20 mM Tris-HCl (pH 7.5), 150 mM NaCl, 5 mM MgCl_2_, 1 mM DTT, 100 µg/ml cycloheximide, and 1% Triton X-100) was added. Tissues were lysed by pipetting and these lysates were incubated with 15 U of TURBO DNase (Invitrogen) for 10 min on ice. Supernatants were collected after centrifugation at 20,000g for 10 min at 4 °C. The Qubit RNA HS Assay Kit (Thermo Fisher Scientific) was used to measure the RNA concentration. Supernatants containing 400 ng RNA was treated with 0.8 U of RNase I for 45 min at 25 °C. Then, same process was performed as previously described ^53^. To remove rRNAs from the total RNA, the Ribo-Zero Gold rRNA Removal Kit (Human/Mouse/Rat) (Illumina) was used.

### Ribo-seq and RNA-seq analysis

Sequence data from RNA-seq and ribosome profiling were trimmed with Trim Galore! (--phred33 -q 30 --length 35) (https://www.bioinformatics.babraham.ac.uk/projects/trim_galore/). Cutadapt (https://cutadapt.readthedocs.io/en/stable/guide.html) was used to remove universal adaptors and linker sequences. Reads were then mapped onto mouse genome mm10 using hisat2 (2.1.0) ^54^ and rRNA was removed from mapped reads. Duplicates were marked and removed with Picard (https://broadinstitute.github.io/picard/). For Ribo-seq, Ribodiff ^55^ was used to count reads and genes which have more than 10 counts in ribo-seq (11370 genes) were used to detect genes with different translational efficiency. For RNA-seq, stringtie (1.3.6) ^56^ was used to identify differentially expressed genes between experiments and the result was further analyzed by TCC (1.22.1) ^57^.

### Chromatin immunoprecipitation (ChIP) analysis

Cells were either FACS-sorted from E11, E12 and E14 *Hes1-d2-EGFP*^*TG/+*^ mice or from E14 cortices of control (*P53*^*-/-*^ *Emx1*^*Cre/+*^, *Fbl*^*flox/+*^*P53*^*-/-*^ and *Fbl*^*flox/flox*^*P53*^*-/-*^), heterozygotes (*Fbl*^*flox/+*^*P53*^*-/-*^ *Emx1*^*Cre/+*^) and DKO (*Fbl*^*flox/flox*^*P53*^*-/-*^*Emx1*^*Cre/+*^). The number of cells was counted as described above. These cells were fixed with 0.25% PFA in PBS for 10 min at RT, washed with 0.1 M glycine in PBS for three times. These cells were collected after centrifugation at 1500 g for 5 min. After remove of supernatant, these cells were resuspended with ChIP buffer (10 mM Tris-HCl pH 8.0, 200 mM KCl, 1mM CaCl_2_, 0.5% NP40) at the concentration of 1×10^6^ cells/ml. After a brief sonication (TOMY HandySonic, 10 s, level 10), micrococcal nuclease (Worthington) was added at the concentration of 50 U/ml. The mixer was incubated at 37 °C for 20 min. EDTA (final concentration: 10mM) was added to stop MNase reaction. The lysates were collected after centrifugation at 15000 g for 5 min and supernatants were removed. The lysates were resuspended with RIPA buffer (50 mM Tris-HCl pH 8.0, 150 mM NaCl, 2mM EDTA, 1% NP40, 0.5% Sodium Deoxycholate, 0.1% SDS). After sonication for three times (10 s, level 10), lysates were centrifuged at 15,000 g for 5 min and supernatants were collected. For each ChIP experiment, 100 µL lysate was used. 25 µL Dynabeads-Anti Rabbit or Mouse IgG (Invitrogen) were washed with 500 µL ChIP buffer. Beads were incubated with 1 µL of primary antibody H3K27me3 (rabbit monoclonal,Cell Signaling Technology, #9733) and H3K4me3 (monocle, wako, 307-34813) in 300 µL blocking buffer (5 mg/ml BSA, 0.5% NP40, 0.1% Tween20 in PBS) at 4 °C with gentle rotation overnight. After wash with ChIP buffer three times, those beads were mixed with 400 µL blocking buffer and 100 µL ChIP lysate and incubated for 1 h at 4 °C. These beads were washed five time with low salt wash buffer (20 mM Tris-HCl pH 8.0, 150 mM NaCl, 2mM EDTA, 1% Triton-×100 and 0.1% SDS) and high salt wash buffer (20 mM Tris-HCl pH 8.0, 500 mM NaCl, 2 mM EDTA, 1% Triton-×100 and 0.1% SDS), respectively, followed by the release of chromatin by incubation of these beads with 200 µL elution buffer (50 mM Tris-HCl pH 8.0, 10 mM EDTA and 1% SDS) for 30 min at 65 °C. The supernatant was removed into a new tube and was incubated at 65 °C for 4 h. 1 µL RNase A (Sigma) was added and incubate at 37 °C for 10 min to degrade RNA. The supernatant was incubated at 55 °C overnight with 5 µL proteinase K (Roche). Genomic DNA was extracted with phenol:chloroform extraction.

### Bioinformatics analysis of ChIP data

Sequenced reads with poor quality were trimmed with Trim Galore! (--phred33 -q 30 --length 35) (https://www.bioinformatics.babraham.ac.uk/projects/trim_galore/). Reads were mapped onto mouse genome mm10 using bowtie ^58^ with the parameter –m1 --best --strata. Mapped sam files were transferred into bam files and were sorted with samtools (1.5) ^59^. Duplicates were marked and removed with Picard (http://broadinstitute.github.io/picard/). Peaks of ChIP-seq were called using MACS2 (2.1.1) ^60^. Q-value to cutoff H3K4me3 peaks was set at 0.01. For call peaks of H3K27me3 --broad function was used and q-value was set at 0.01 and 0.05 to cutoff narrow/strong regions or broad/weak regions, respectively. For each sample, specified input was used as control.

Deeptools (3.2.1) ^61^ was used to calculation the correlation of each data set. The alignment files were binned using multiBamSummary function with default setting and pearson correlation was calculated using plotCorrelation function. To confirm the quality of our ChIP-seq data, we also compared our H3K4me3 and H3K27me3 ChIP-seq using E14 NSC with the published H3K4me3 (ENCSR172XOZ) and H3K27me3 (ENCSR831YAX) ChIP-seq data using E14 forebrain (https://www.encodeproject.org/).

ChromHMM (1.14) ^62^ was used to evaluate state transition between different stages. The alignment files of H3K4me3 and H3K27me3 in each stage were binned into 200-bp bins using BinarizeBam. Then, we established the model with 4 emission states (H3K4me3-only, H3K27me3-only, bivalent and none) and trained with binned data using LearnModel command. These segmentation files with state information was used to plot alluvial plotting.

DiffBind package (2.10.0) ^63^ in R was used to find peaks with different intensity between samples and different stages and to visualize data with PCA. To do so, overlapping peaks among each samples were isolated and sequenced reads on these consensus peaks were counted using dba.count function. As a result, a matrix in which each column indicates a consensus peak and each row indicates the normalized reads counting was produced. The matrix was used to plot PCA using ggplot function in R. The peaks with different intensity was calculated using dba.contrast and dba.analyze function. MA plotting was draw by dba.plotMA function. Circos plot was generated by Circos tool ^64^ using H3K27me3 peaks showing different intensity between E11 and E14 NSCs.

### Library preparation and sequencing

Library preparation for RNA-sequencing (RNA-seq) was performed as described ^65^ using the TruSeq Stranded mRNA Library Prep Kit (Illumina) and 100-110 ng of total RNA. Library preparation for ChIP-sequencing (ChIP-seq) was performed as described ^66^ using the KAPA LTP Library Preparation Kit (KAPA Biosystems) and 2 ng of input DNA, or the entire amount of the ChIP DNA obtained. KAPA Real-Time Library Amplification Kit (KAPA Biosystems) was used in conjunction with the library preparation kits described above to minimize the number of PCR cycles for library amplification. Library preparation for single-cell RNA-seq (scRNA-seq) was performed following the standard protocol of the 10x Genomics Chromium Single Cell 3′ v2 Kit (10x Genomics). Sequencing was performed in HiSeq1500 (Illumina) with the HiSeq SR Rapid Cluster Kit v2 (Illumina), the HiSeq PE Rapid Cluster Kit v2 (Illumina), or the TruSeq PE Cluster Kit v3-cBot-HS, to obtain single-end 80 nt reads for RNA-seq and ChIP-seq libraries, or paired-end 26 nt (Read 1)-98 nt (Read 2) reads for scRNA-seq libraries. Total reads of each sample were reviewed in Extended Data Table 6.

### Primary cell culture

For knockdown experiment, cells from E11 dorsal cortices were counted and resuspended with buffer R (Neon™ transfer system, Invitrogen) in the concentration of 8×10^6^ cells/ml. Resuspended cells (100 µL) were mixed with 160 µM control or *Fbl* siRNA. Electroporation was performed with Neon (Neon™ transfer system, Invitrogen) at condition of 1600 voltage, 20 width and 1 pulse. These transfected cells were mixed with 2 ml culture medium (20 ng/ml human basic FGF (Peprotech), 1XB27 RA-(Gibco), in Dulbecco’s Modified Eagle Medium (DMEM) F12+GlutaMax (Gibco)) and distributed into 4 well or 24 well-plates (500 µl/well) and cultured at 37 °C. For inhibitors treatment, 2×10^5^ cells/well were culture in the DMSO, 2.5 µM GSK-J4 (Sigma, SML0701), 2.5 µM GSK-343 (Sigma, SML0766) or 2.5 µM GSK-J4 and 2.5 µM GSK-343 for 3 days.

For test cap-independent translational activity, the reporter (4 ug) with 160 µM control or *Fbl* siRNA was transfected into 100 µL of resuspended cells from E11 dorsal cortices as described above. After 2 days, these cells were washed with PBS and treated with 0.05% Trypsin/HBSS for 10 min. Then, these cells were washed with 0.375% BSA/HBSS and harvested for sorting.

For cell cycle analysis, *Hes5-d2-EGFP* plasmid (2 µg) with 160 µM control or Fbl siRNA was transfected into resuspended cells from E11 dorsal cortices as mentioned above. After 2 days, cells were harvested for sorting as described above. GFP positive NSCs were sorting into 1 mL of 0.375% BSA/PBS with SH800 (SONY). Then, ice-cold 100% ethanol (3 ml) were added into these sorted cells with gently vortex. These cells were fixed at −30 °C for 1 h, followed by centrifugation at 1500 g for 5 min. After remove of supernatant, 700 µL of staining solution (50 µg/ml propidium iodide (nacalai tesque) and 100 µg/ml RNase A in 1% BSA/PBS) was added and mixed well by pipetting. Sorting of cells was performed by SH800 according with the manufacturer′s protocol.

### O-propargyl-puromycin (OPP) visualization

Cells from E14 dorsal cortices with different genotypes were isolated and cultured for one day as described above. Two µL of OPP reagent was added into medium and incubated for 30 min. These cells were fixed with 1% PFA overnight and OPP signal was detected using Click-iT™ Plus OPP Alexa Fluor™ 488 Protein Synthesis Assay Kit (Invitrogen) as the manufacturer′s protocol. To distinguish NSCs, immunostaining was performed with Sox2 antibody. OPP signal was measured in Sox2 positive cells using CellProfiler (2.1.1) ^67^.

### Protein stability measurement

Cells from E11 dorsal cortices were treated with 100 µg/ml cycloheximide for 0.5, 1 and 2 h followed by collection for western blot. As control, cells without treatment were also collected.

### Knockout of 5′UTR of *Ezh2* and *Kdm6b*

Two µg of pCAG-EGFP plasmid with 1 µg of gRNA plasmid for 5′UTR of *Ezh2* and *Kdm6b* and 1 µg of pCAX-Cas9 was transfected into cells from E11 cortices with NEON and these cells were cultured as mentioned above. Two days after transfection, GFP positive cells were sorting with the SH800 (SONY). These cells were used for reverse transcription (RT)-quantitative (Q) PCR, western blot and genotyping. For cells used for genotyping, 5µL protein kinase K was added and incubated for 1 h. After the treatment at 98°C for 5 min, genotyping was performed used following primer: *Ezh2* forward:5′-GAATTCTGCAGTCGACGCTTGATAGTGCTGGGGG-3′ *Ezh2* reverse: 5′-CCGCGGTACCGTCGACGCCGAAGACTGGCCAGGC-3′ *Kdm6b* forward:5′-GAATTCTGCAGTCGAGGCCTGGGTGCTGGATTTG −3′ *Kdm6b* reverse: 5′-CCGCGGTACCGTCGATCAGCATCAAGAGCCCCTAG-3′. PCR products of these primers was cloned into pCAG plasmid cut with SalI and sequencing. For the cells used for RT-QPCR, total RNA was extracted by RNeasy Mini Kits (Qiagen) and genomic DNA was removed by the treatment of DNase (Qiagen). RNA (30 ng) was used for synthesis of cDNA with PrimerScript RT reagent kit with gDNA eraser (TAKARA BIO). QPCR was performed with TB Green Premix Ex-taq II (TAKARA BIO). *Gapdh* was used as internal reference to normalize the dosage of *Ehz2* and *Kdm6b* mRNA level, following primers were used for QPCR: *Ezh2* forward: 5′-GAGCGTATAAAGACACCACCTAAAC-3′; *Ezh2* reverse: 5′-CTCTGTCACTGTCTGTATCCTTTG-3′. *Kdm6b* forward: 5′-CCCCCATTTCAGCTGACTAA-3′; *Kdm6b* reverse: 5′-CTGGACCAAGGGGTGTGTT-3′; *Gapdh* forward: 5′-ATGAATACGGCTACAGCAACAGG-3′; *Gapdh* reverse: 5′-CTCTTGCTCAGTGTCCTTGCTG-3′. The cells used for western blot were treated as described above.

### Knock out of *Fbl* and *Trp53* by CRISPER/Cas9 system

The gRNA plasmid together with pCAX-Cas9 were transfected into cells from E10 dorsal cortices with NEON as described above. To label knockout cells, EGFP was simultaneously knocked into *beta-actin* locus as described before ^49^. For 2×10^5^ cells, 0.5 ug of each plasmid was used. For clone analysis, 10,000 electroporated cells were mixed with 190,000 untreated cells and cultured for 4 days. Culture medium was changed every 2 days. To knockout of *Fbl* and *Trp53 in utero*, 0.3 ug/µL gRNA sets and pCAX-Cas9 were electroporated into dorsal brains at E11. *In utero* electroporation in mice was reported previously^68,69^.

### Calculation of rRNA level

Dorsal cortices were removed from E14 mice with different genotypes. RNA extraction was performed as described above. To examine rRNA level, we used the method described before with modification^70^. Briefly, 10 ng RNA was mixed with 1 µM random primer and 1 mM dNTP (high condition) or 0.004 mM dNTP (low high) and incubated 65°C for 5 min. Then, 5× First Stand buffer (invitrogen), 10× SuperScript® III Reverse Transcriptase (invitrogen), 0.005 mM DTT and 20× RNase inhibitor (TAKARA BIO) was added and the mix was incubated at 50 °C for 1 h followed by 70 °C for 15 min. QPCR was then performed as described above to determine the dosage of amplicon of each primer at different conditions. Following primer sets were used: 28S_1673 forward: 5′-CTAGTGGGCCACTTTTGGTA-′3; 28S_1673 reverse: 5′-TTCATCCCGCAGCGCCAGTT-′3; 28S_2614 forward: 5′-TAGGTAAGGGAAGTCGGCAA-′3; 28S_2614 reverse: 5′-CCTTATCCCGAAGTTACGGA-′3; 28S_3441 forward: 5′-ATGACTCTCTTAAGGTAGCC-′3; 28S_3441 reverse: 5′-TCACTAATTAGATGACGAGG-′3; 28S_4223 forward: 5′-GGTTAGTTTTACCCTACTGA-′3; 28S_4223 reverse: 5′-GATTACCATGGCAACAACAC-′3; 28S_4242 forward: 5′-TGATGTGTTGTTGCCATGGT-′3; 28S_4242 reverse: 5′-GTTCCTCTCGTACTGAGCAG-′3; 28S_3958 forward: 5′-CTCGCTTGATCTTGATTTTC-′3; 28S_3958 reverse: 5′-CGCTTTCACGGTCTGTATTC-′3; 28S_4188 forward: 5′-TAGGGAACGTGAGCTGGGTTTAGA-′3; 28S_4188 reverse: 5′-GTAAAACTAACCTGTCTCACGACG-′3; Methylation level of rRNA was calculated as dosage of amplicon at low condition/ dosage of amplicon at high condition.

### Motif find and calculation of minimum free energy

To find the motif of mRNAs that downregulate or upregulate their translational efficiency after knockout of *Fbl*, 5′ UTR sequence of selected genes were initially extracted from Table Browser (UCSC) http://genome.ucsc.edu/cgi-bin/hgTables. Next, HOMER was used to extract enriched motif by findMotifsGenome.pl function with size=200. For each 5′ UTR, minimum free energy was calculated every 50bp by ViennaRNA Package 2.4.14 ^71^ with RNALfold L50 –g and the minimum value was used.

### Statistics

Multiple comparisons for cell counting and 5′UTR knockout experiments were done using one-way ANOVA followed by a Tukey′s HSD. Student t-test was used to test cap-independent translational activity after knockdown of *Fbl*. To test whether methylation level on rRNA are reduced in DKO comparing with control cells, one-sided wilcoxon signed rank test was used. Kruskal-Wallis test with Dunn′s multiple-comparison test was used to test different PC contribution of genes and global protein level among different genotypes of mice. Wilcoxon signed rank test was used to test MEF of different gene groups. R, RStudio, and Excel (Microsoft) were used for these analyses.

## Acknowledgments

We thank A. Shitamukai, I. Fujita, Y. Tsunekawa, D. Konno and all members in Matsuzaki laboratory for their discussion and technologic advices. We thank C. Tanegashima, K. Tatsumi, and O. Nishimura of the Laboratory for Phyloinformatics, RIKEN Center for Biosystems Dynamics Research (BDR) for NGS library preparation, sequencing, and assistance on data production and analysis. This work was supported by the Japan Society for the Promotion of Science (JSPS) Grants-in-Aid for Scientific Research (KAKENHI) grand no. 18K14722 to Q.W., Scientific Research on Innovative Areas 17H05779 and 19H04791, and RIKEN funds to F.M. Q.W. was also supported by JSPS Postdoctoral Fellowship and RIKEN Special Postdoctoral Researcher Program.

## Author contributions

F.M. supervised the project. F.M. and Q.W. designed experiments and wrote the manuscript. Q.W. carried out experiments and performed the bioinformatics analysis of data. Y.S. and S.I developed the methods for ribosome profiling using dorsal cortices and helped Q.W. in the analysis of ribosome profiling data. T.A and H.K. generated *Fbl* conditional knockout mice. T.S. did *in utero* electroporation. A.O helped Q.W. for sequencing, QPCR, and purification of plasmids.

## Data and materials availability

All raw and proceeded data for scRNA, ChIP-seq and Ribo-seq are available at the DNA Data Bank of Japan with accession number DRA009567, DRA009568, DRA009569 and DRA009729, and E-GEAD-348, E-GEAD-349, E-GEAD-350 and E-GEAD-351. Code used in this study including WGCNA, Cytoscape (3.7.2), Seurat (2.3.4), monocle (2.10.1), Trim Galore! (0.4.2), hisat2 (2.1.0), Ribodiff (0.2.1), bowtie (1.2.1.1), Picard, Samtools (1.5), MACS2 (2.1.1), ngsplot (2.61), IGV (2.4.3), Deeptools (3.2.1), ChromHMM (1.14), DiffBind (2.10.0), CellProfiler (2.1.1), ViennaRNA (2.4.14), stringtie (1.3.6) and TCC 1.22.1 are available from indicated manual or references.

---

**Extended Data Fig.1.**
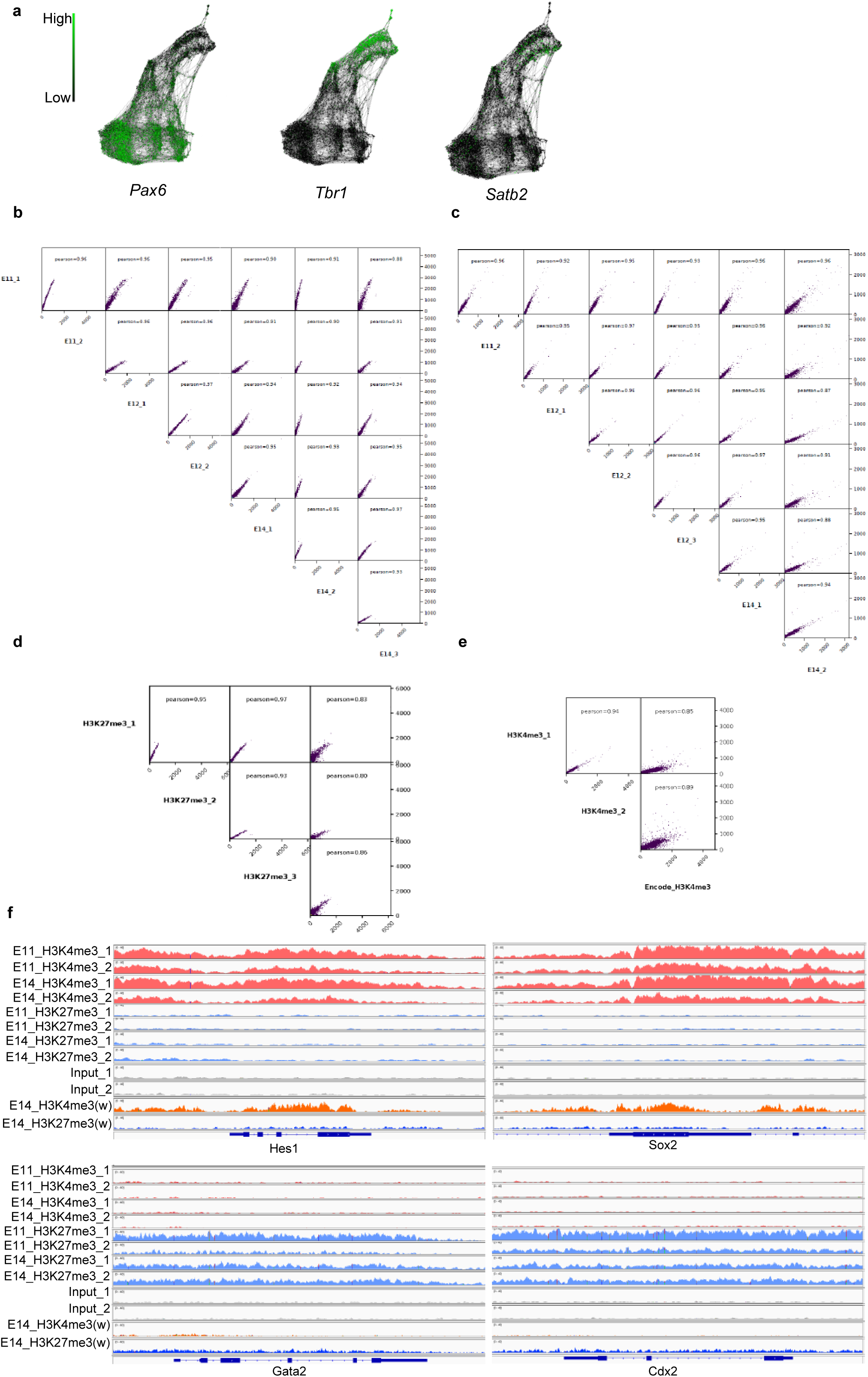
Quality check of single cell RNA-seq and ChIP-seq experiments. **a**, SPRING graphs indicating the expression pattern of a NSC marker: *Pax6*, a deep layer marker: *Tbr1* and an upper layer marker: *Satb2*, respectively. **b,c**, Sample correlation of ChIP-seq experiments using H3K27me3 (b) and H3K4me3 antibodies (c among E11, E12, and E14 NSCs. **d,e**, Sample correlation of ChIP-seq experiments using H3K27me3 (d) and H3K4me3 antibodies (e) between our data from E14 NSCs and published data from E14 whole brains. **f**, Genome browser view of ChIP-seq density of the indicated genes.

**Extended Data Fig.2.**
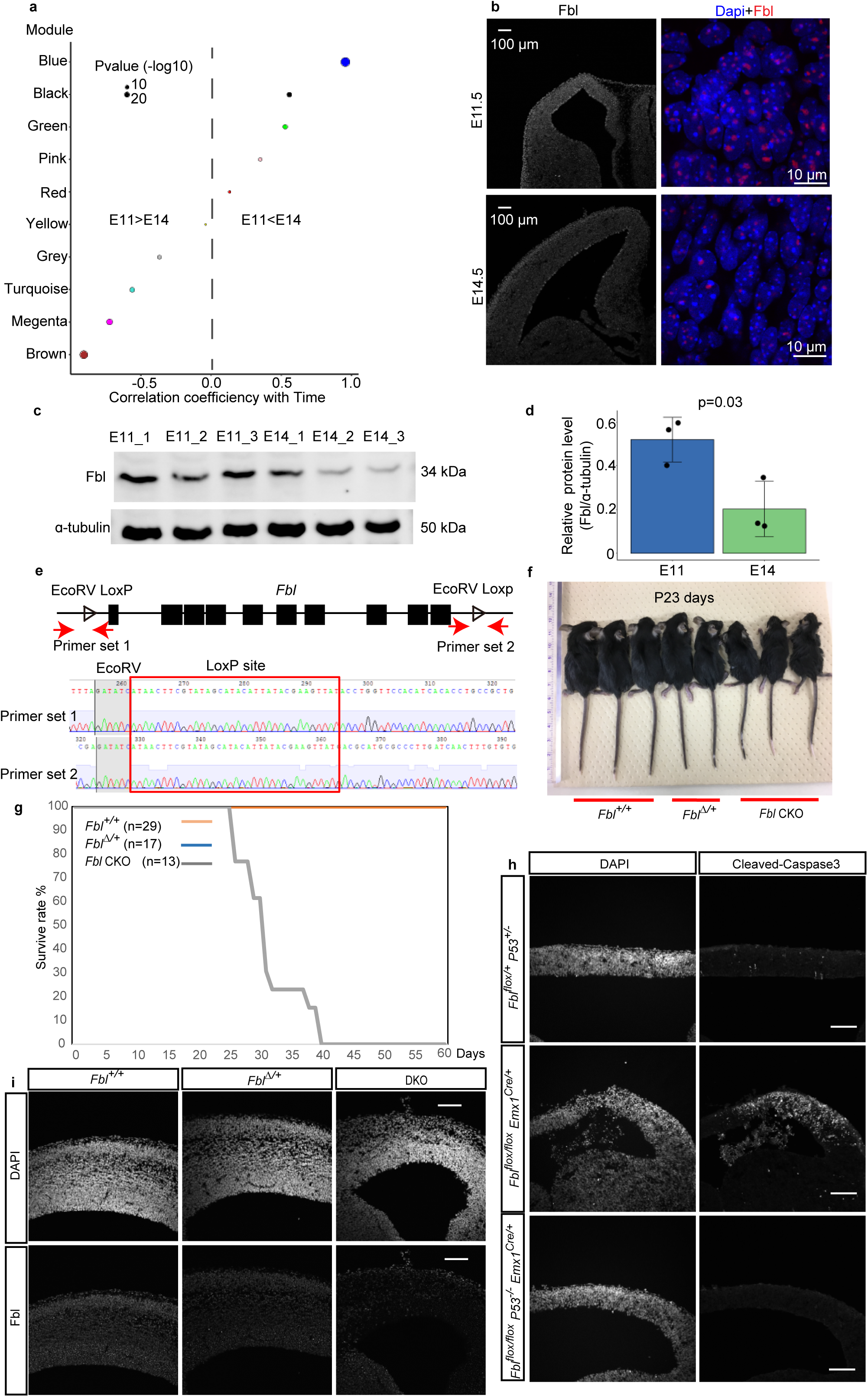
Identification of Fbl as a condidate to promote the developemental clock in NSCs. **a**, WGCNA gene dendrogram classifies E11 and E14 NSCs into different modules and correlation coefficient of each module with time (E11 and E14). **b**, Representative image of E11 and E14 brain sections stained for Fbl. Scale bar: 10 or 100 µm **c**, Western blotof Fbl and α-tubulin using isolated Hes1+ NSCs. **d**, Quantitative analysis of Fbl protein levels showing reduced Fbl at E14 (Student t-test; data are presented as mean±s.d.). **e**, Schematics of CRISPR-CAS9-dependent knock-in of loxP sites flanking Fbl. Confirmation of loxP sites by sequencing analysis. **f**, Appearance of Fbl^+/+^, Fbl^Δ/+^, and Fbl CKO pups at 23 days postnatal. **g**, Survival rate of Fbl^+/+^, Fbl^Δ/+^, and Fbl CKO pups. **h**, Representative image of E12 brain section stained for cleaved-caspase 3 antibody. Scale bar: 100 µm. **i**, Representative image of E14 brain section stained for Fbl. Scale bar: 100 µm.

**Extended Data Fig.3.**
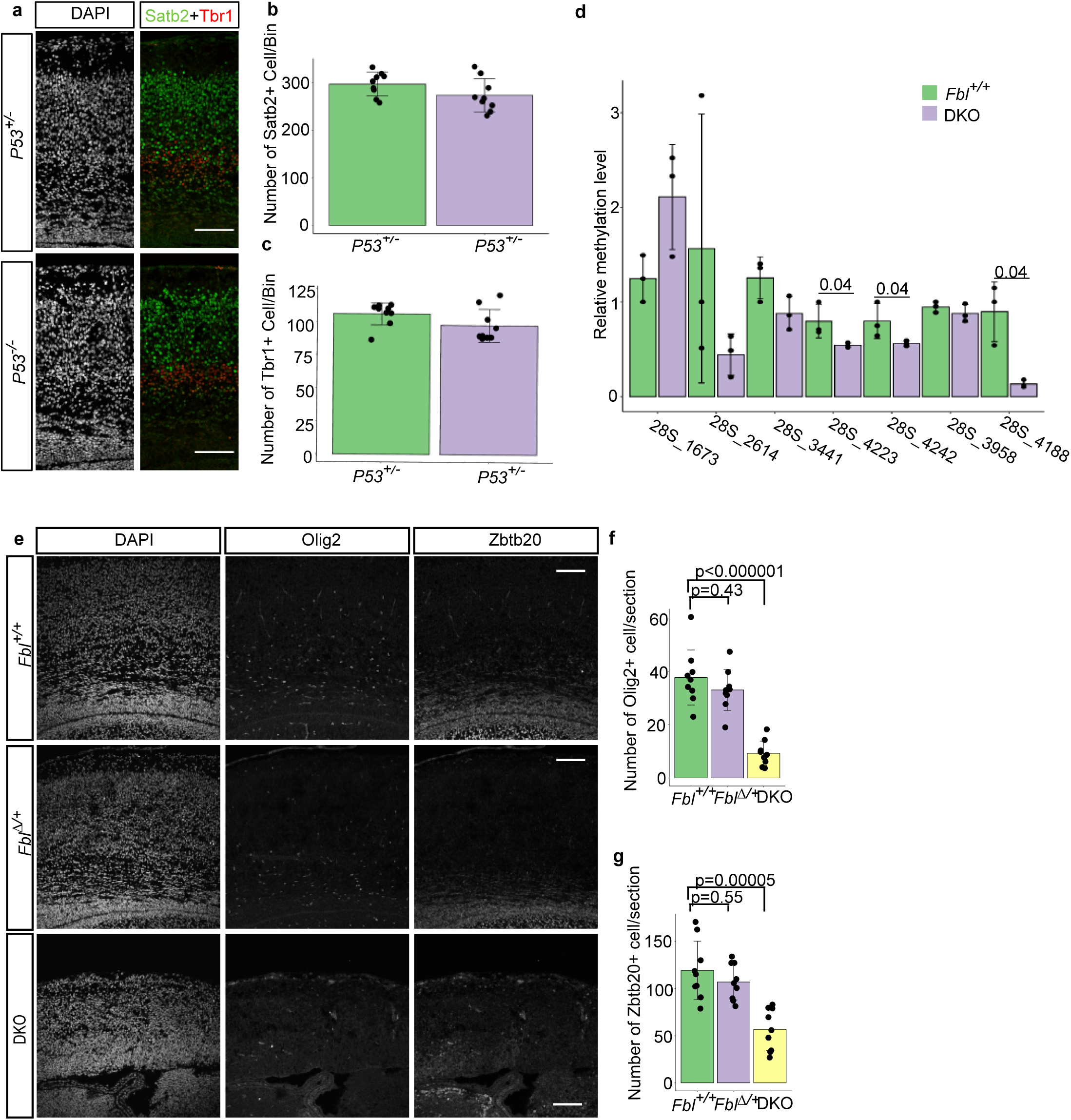
Deletion of *Trp53* does not affect neurogenesis, while *Fbl* knockout affects methylation of rRNA, oligogenesis and astrocytogenesis. **a**, Representative image of *Trp53*^*+/+*^ and *Trp53*^*-/-*^ brain sections stained for Satb2 and Tbr1. **b,c**, Cell number quantification on sections based on immunostaining with the indicated markers (n=3 mice per genotype, n=3 sections per mouse). **d**, Methylation level at the indicated sites in *Fbl*^*+/+*^ and DKO NSCs (one-sided Wilcoxon signed rank test; data are presented as mean±s.d). **e**, Representative image of E17 brain section stained for Oligo2 and Zbtb20. **f,g**, Quantification of cell number on sections basing on immunostaining with the indicated markers (n=3 mice per genotype, n=3 sections per mouse; one-way ANOVA followed by Tukey’s post-hoc tests; data are presented as mean±s.d. of n=9 sections). Scale bar: 100 µm.

**Extended Data Fig.4.**
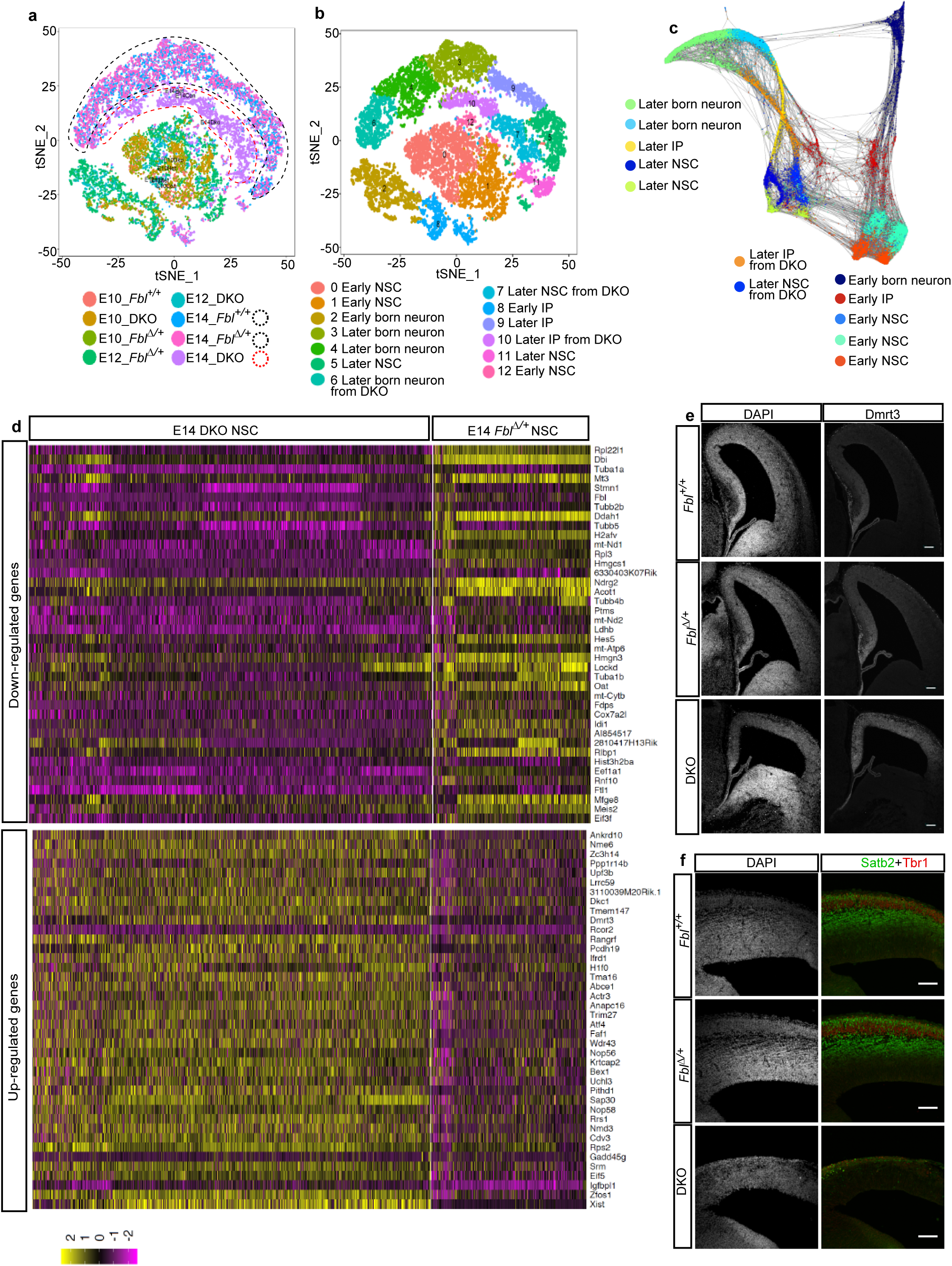
Single-cell transcriptome analysis of temporal pattern in NSCs. **a,b**, Scatterplot of single-cell transcriptome after t-stochastic neighbour embedding (t-SNE), coloured by genotypes from different stages (a) or different cell types (b). In (a), the separation of E14 DKO brain cells from E14 *Fbl*^*+/+*^ or *Fbl*^*Δ/+*^ brain cells is emphasized by the dotted red and black circles, respectively. **c**, SPRING graph of single cells colored by cell type identified by the expression of marker genes. **d**, Heatmap of the top 40 differentially expressed genes in the comparison of E14 DKO and *Fbl*^*Δ/+*^ NSCs, ranked by fold change. **e,f**, Representative images of E14 brain sections stained for Dmrt3 (e) and Satb2/Tbr1 (f). Staining was performed in three different biological samples for each genotype. Scale bar: 100 µm.

**Extended Data Fig.5.**
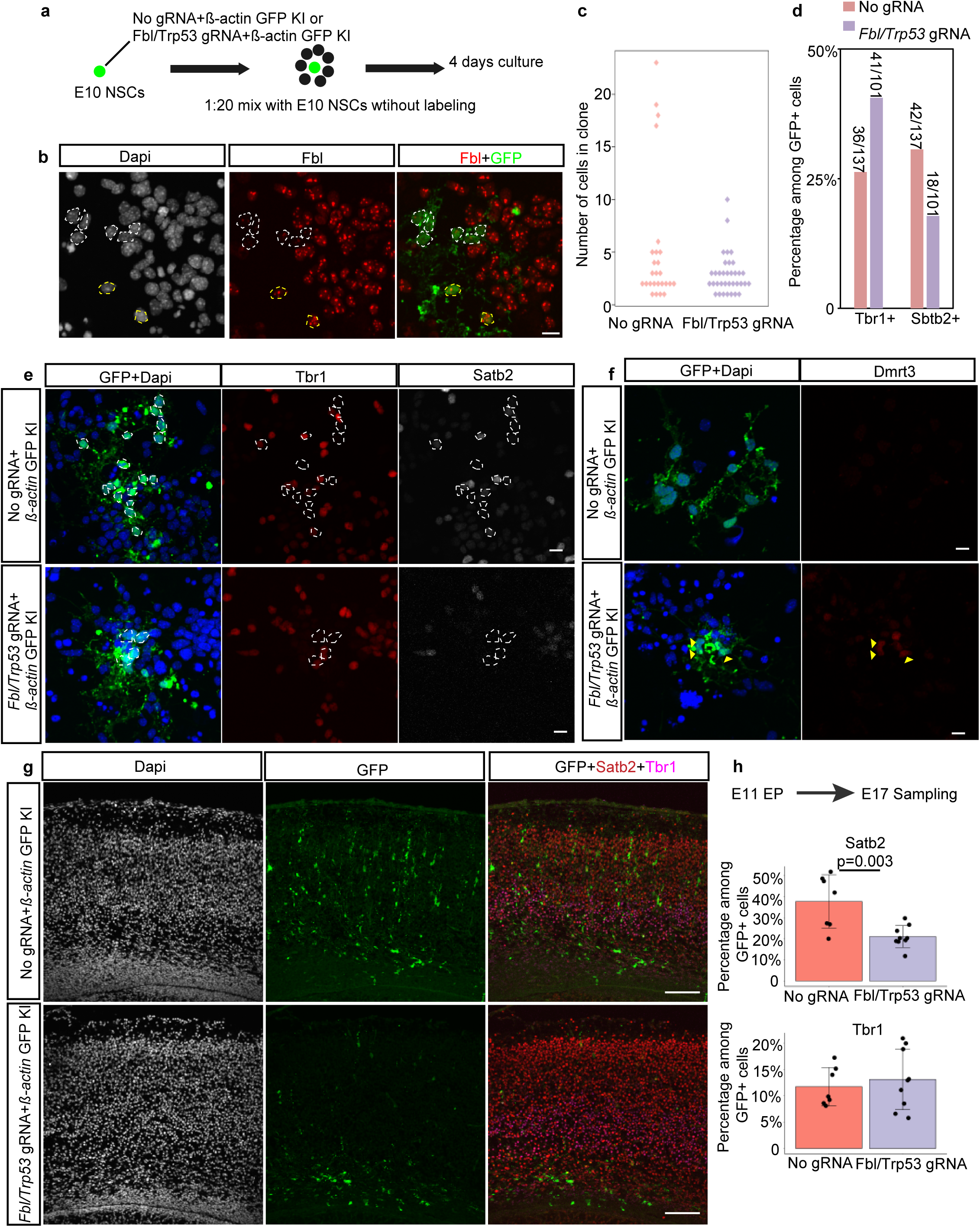
Fbl works intrinsically for temporal pattern transition. **a**, Schematics of the experimental design for clonal analysis of *Fbl/Trp53*-null NSCs. Cells from dorsal brains were electroporated at E10 with/without *Fbl* and *Trp53* gRNA for deletion and with β-actin gRNA for labelling. **b**, Representative image of cultured cells stained for Fbl showing deletion of Fbl in some GFP-positive cells (n=39/88). White and yellow circles indicate Fbl-negative and positive cells, respectively. Scale bar: 10 µm **c**, Clone size analysis of normal and *Fbl/Trp53*-null NSCs after 4 days of culture. (Clone number n=26 and n=35 for control and knockout, respectively, from two independent experiments) **d**, Percentage of Tbr1- and Satb2-positive cells among GFP-positives based on staining. The number of counted cells is shown. **e**, Representative image of cultured cells stained for GFP/Dapi, Tbr1, and Brn2. Scale bar: 10 µm. **f**, Representative image of cultured cells stained for GFP/Dapi and Dmrt3, showing more *Fbl/Trp53*-null cells (11/39) expressing Dmrt3 than control cells (2/34), which was also observed in *Fbl* DKO brains in Figure S4E. Scale bar: 10 µm. **g**, Representative image of E17 brain section stained for GFP, Tbr1, and Satb2. Dorsal brains were electroporated at E11 with/without *Fbl* and *Trp53* gRNA for deletion and β-actin gRNA for labelling. Scale bar: 100 µm. **h**, Quantification of Satb2- and Tbr1-positive cells on sections based on immunostaining (n=3 mice per genotype, n=2-3 sections per mouse; Student’s t-test; data are presented as mean±s.d. of counted sections).

**Extended Data Fig.6.**
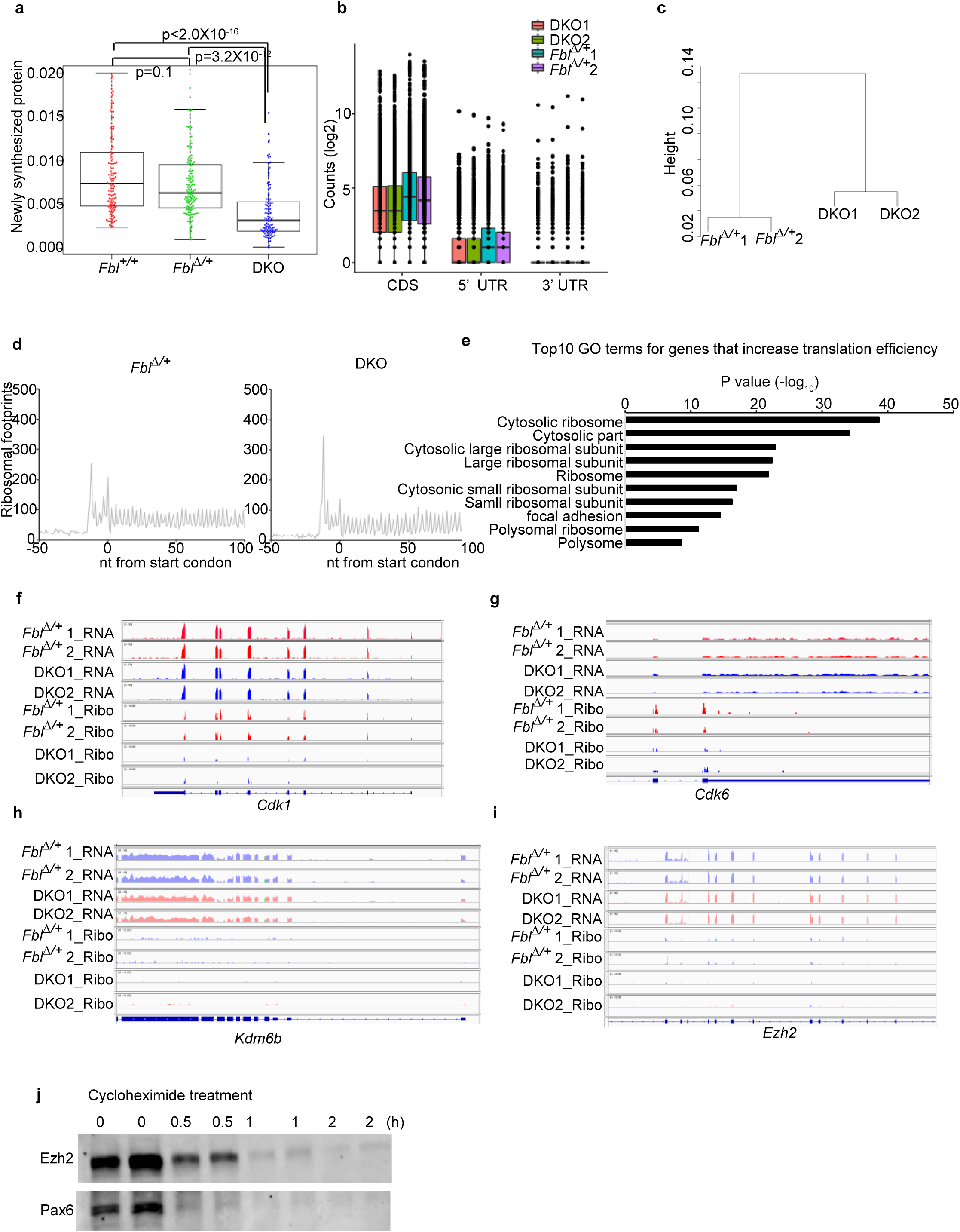
Assessment of translational efficiency (TE) dependent on Fbl via ribosome profiling. **a**, Quantification of O-propargyl-puromycin (OPP) incorporation in NSCs (Kruskal-Wallis test and Dunn’s test with Bonferroni correction; Fbl^+/+^: n=135; Fbl^Δ/+^: n=148; DKO: n=106). **b**, Counts of ribosome footprints mapping on different mRNA regions. **c**, Hierarchical clustering of ribosome profiling data. **d**, Metagene analysis of the 5’ end of footprints showing three-nucleotide periodicity of the ribosomal footprint reads. **e**, Top 10 GO terms of transcripts whose TE increased after Fbl knockout. **f-i**, Genome browser view of RNA-seq and Ribo-seq density of the indicated genes. **j**, Western blot analysis of Ezh2 and Pax6 after treatment with 100 µg/ml cycloheximide for the indicated time, showing that Pax6 was not more stable than Ezh2 (n=2).

**Extended Data Fig.7.**
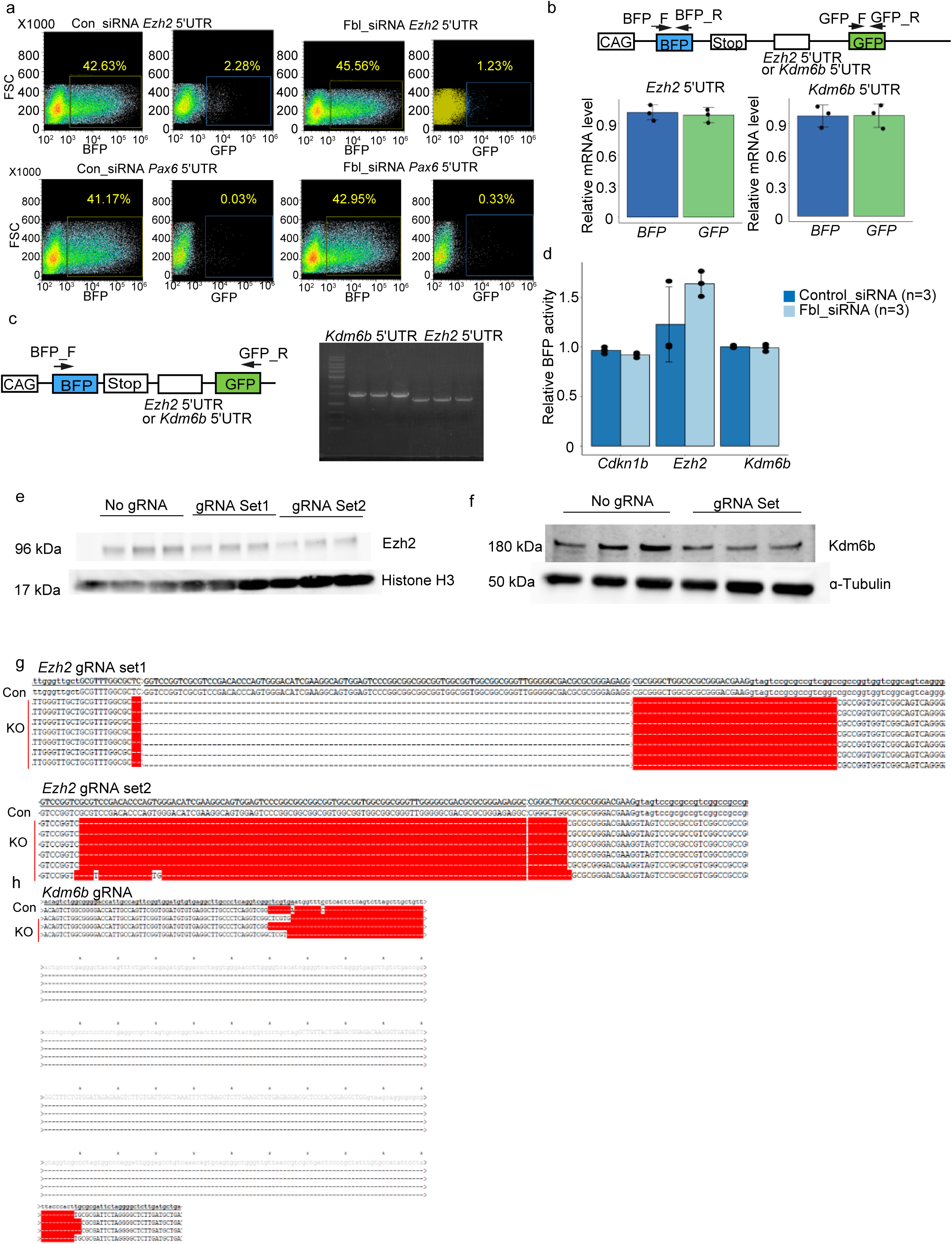
Cap-independent translational activity in the 5′UTR of Fbl target mRNAs. **a**, Representative plot of sorting GFP and BFP populations using the 5′UTR of Pax6 and Ezh2. **b**, qPCR of BFP and GFP from transfected cells indicates the same expression level of BFP and GFP (data are presented as mean±s.d., n=3). **c**, RT–PCR using primers in BFP and GFP confirms the absence of cryptic splices in the 5′UTR of Ezh2 and Kdm6b (n=3 each). **d**, Quantitative analysis of BFP signal after Fbl knockdown (data are presented as mean±s.d., n=3) **e,f**, Detection of Ezh2 and Kdm6b protein by western blot after knockout of the 5′UTR of these genes. **g,h**, Confirmation of the deletion of the 5′UTR of Ezh2 (g) and Kdm6b (h) by sequencing.

